# Optogenetic stimulation of primate V1 reveals local laminar and large-scale cortical networks related to perceptual phosphenes

**DOI:** 10.1101/2021.06.01.446505

**Authors:** Michael Ortiz-Rios, Beshoy Agayby, Fabien Balezeau, Marcus Haag, Samy Rima, Michael C. Schmid

## Abstract

Developing optogenetics in non-human primates (NHPs) is essential for translating its successful implementation in rodents to clinical applications in humans. However, information about how optogenetics influences the primate cortex remains limited. Here, we evaluate how optogenetic stimulation of the primate primary visual cortex (V1) affects local and large-scale network activation concerned with visual perception. To this end we injected an optogenetic construct (AAV9-hSyn-ChR2-eYFP) into the V1 cortex of four macaque monkeys (*macaca mulatta*) and measured the effects of optogenetic V1 stimulation using functional magnetic resonance imaging (fMRI), laminar electrophysiology, and behavioural assessment. In three macaques, blood-oxygen-dependent (BOLD) fMRI activity could be reliably elicited with optogenetic stimulation in V1 and several connected extrastriate brain areas, including V2/V3, motion-sensitive area MT and the frontal-eye-fields (FEF), in particular when pulsed stimulation at 40 Hz was applied. BOLD modulation was associated with consistent neural spiking activity measured in V1 of two macaques. More detailed analysis revealed strongest neuronal activation in layer 4B and infragranular layers, which tightly reflected the histological expression pattern of the optogenetic construct in V1. Driving this visual network proved sufficient to elicit a visual percept (‘phosphene’) in one macaque during a perceptual choice task. Taken together, our findings reveal the laminar and large-cortical activation pattern related to visual phosphene generation and emphasize the need for further improving optogenetic methods in NHPs as a step towards applications in humans.

## Introduction

Recent advances in optogenetic applications in non-human primates (NHPs) are unravelling functional circuit properties with greater neuronal specificity than traditional methods, such as electrical microstimulation and pharmacology, and therefore show great promise in contributing to a refined understanding of neural circuit function in the healthy and the diseased brain (El-Shamayleh and Horwitz, 2019; Galvan et al., 2017; Tremblay et al., 2020). However, optogenetics in NHPs still faces many methodological challenges before its full potential can be unfolded. Crucially, while electrical stimulation is routinely used with great success in research and clinical settings to influence sensory-motor function and cognition, optogenetic manipulation of behaviour in NHP remains challenging (Galvan et al., 2017; Tremblay et al., 2020). Delineating how optogenetic stimulation affects the primate functional circuitry, both at the local and global level is crucial for achieving a deeper understanding of how optogenetic methods may influence behaviour (Ju et al., 2018; Klink et al., 2021). Among several promising approaches, optogenetics has been combined with multi-contact electrophysiology to investigate optogenetic influences on primate cortical circuits with laminar resolution (Klein et al., 2016). Additionally, opto-fMRI has been adapted for NHPs to delineate the large-scale connectivity from optogenetic stimulation (Gerits et al., 2012; Ohayon et al., 2013).

Here we focus on the macaque visual system, and specifically on the primary visual cortex (V1), which has been a long-standing target for delineating the neural basis of conscious vision (Leopold, 2012; Tong, 2003). V1 is the first cortical area known to integrate visual information within its laminar microcircuitry, before relaying visual information into higher-order cortical regions (Callaway, 1998; Casagrande and Kaas, 1994; Felleman and Van Essen, 1991; Hubel and Wiesel, 1968). Its high accessibility for implants combined with a retinotopic organisation that supports high-resolution vision are important reasons why V1 is regularly made the target for the development of methods aimed at restoring visual function in retinal disease (Beauchamp et al., 2020a; Bosking et al., 2017; Tehovnik et al., 2009). In groundbreaking attempts during the 1960s to develop a cortical prosthesis, Brindley & Lewin managed to induce brief precepts of light (“phosphenes”) in a human patient by stimulating parts of its V1 using surface electrode arrays (Brindley and Lewin, 1968). Ever since these initial findings, electrical microstimulation has been further refined, largely also thanks due to the opportunity for investigations in NHP (Chen et al., 2020; Schiller et al., 2011; Tehovnik and Slocum, 2007), where the generation of phosphenes in precise and distinct locations of the visual field could be reported using behavioural testing paradigms that provided the valuable groundwork for testing novel stimulation methodologies such as optogenetics and how they affect visual perception. Similar to other cognitive domains in which optogenetic methods have been tested in NHP with the aim to alter or induce behaviour, it has been difficult to elicit phosphenes from optogenetic V1 stimulation. Recent advancements include elicitation of saccadic eye movements (Jazayeri et al., 2012) and improvement of visual detection (Andrei et al., 2019) from optogenetic stimulation of V1. Similarly, disruption of eye movements and visual detection could be achieved from optogenetic V1 inhibition (De et al., 2020). Critically, however, it remains unclear whether monkeys perceive a phosphene during optogenetic stimulation of V1, which may in part be due to limited knowledge on how V1 optogenetics interfaces with V1’s microcircuitry and large-scale cortical connectivity. To fill this gap we report here from our investigations, in which we first analyzed how optogenetic stimulation results in topographic and layer specific recruitment of V1, before continuing our assessment of large scale activation using opto-fMRI and description of behavioural tests aimed at characterizing the visual percept that results from V1 optogenetic stimulation in NHP.

## Results

### Optogenetic V1 stimulation induces local BOLD and spiking activity

Our first aim was to map the local neural activation induced by the optogenetic stimulation of V1. During a prior surgical procedure, we had injected ∼24 μl of *AVV9-hSyn-hChR2* across multiple depths and sites of opercular V1 (approx. around 5-7° of visual eccentricity retinal representation) (**Fig. 1A**). The injection resulted in an estimated virally transfected area of approx. 12 mm^2^ (see **Table1**). For stimulation, a large-volume LED illuminator was placed epidurally inside an MRI-compatible recording chamber (Ortiz-Rios et al., 2018) implanted over the occipital lobe for chronic access to V1 (**Fig. 1B** and **Supp. Fig. 1B)**. The illuminator consisted of a 1.5 mm diameter optical-fiber with an effective 60° light beam (**Fig. 1C**) that was connected to an external LED producing blue light (451 nm) for effective stimulation of channelrhodopsin (ChR2). The LED stimulation beam allowed us to deliver light to V1 with an effective distance greater than 3.5 mm in depth and 3 - 5 mm laterally along the cortical surface as determined during control experiments in agar (**Supp. Fig. 2A**).

**Figure 1.**
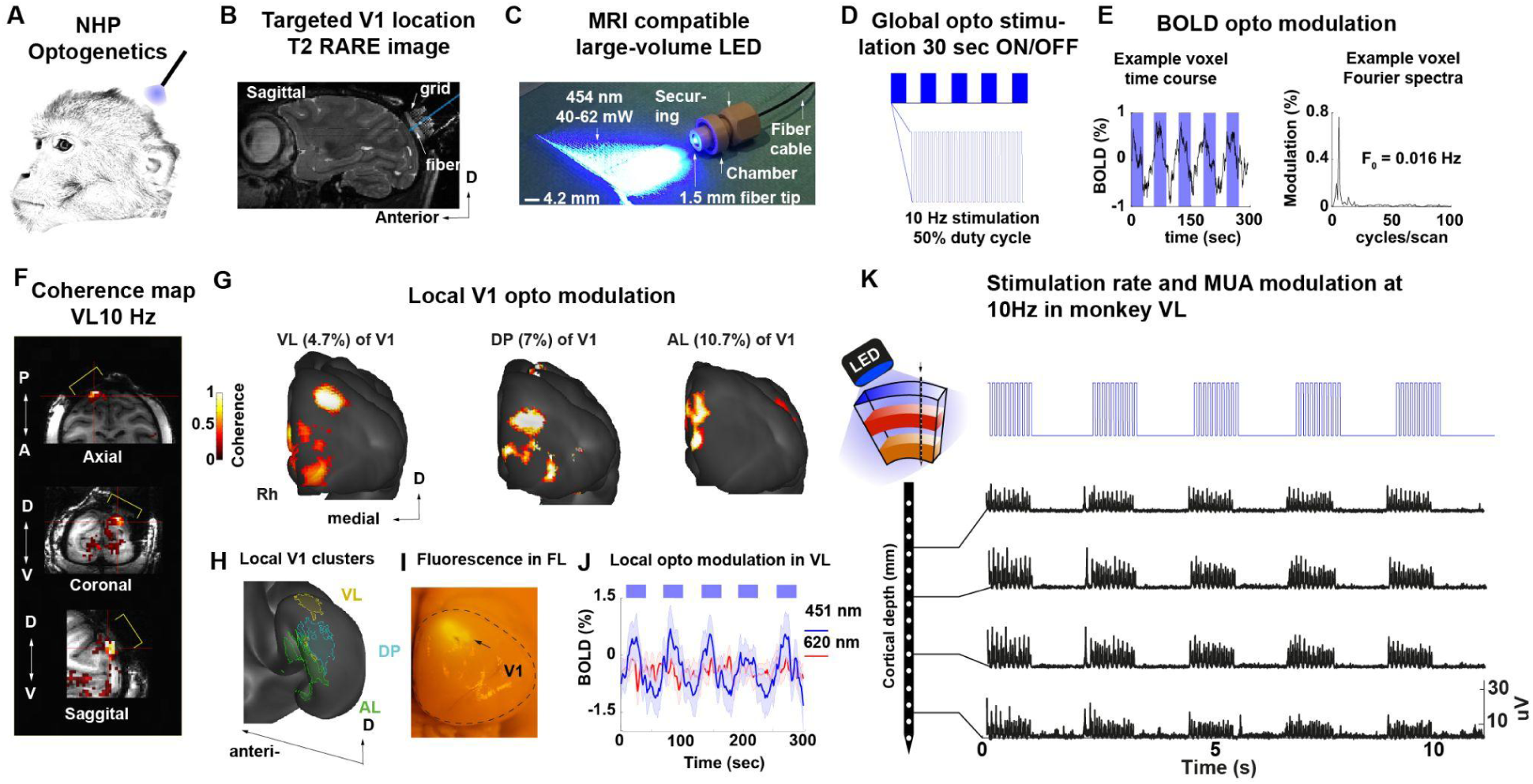
Optogenetic stimulation drives the local BOLD modulation and spiking activity in transfected V1 neurons. **A.** Optogenetic stimulation of V1 in awake NHPs. **B.** T2-RARE sagittal image showing the saline-filled chamber along with the center coordinate of the optical fiber placement on monkey VL. **C**. MRI compatible LED fiber system with a tip diameter of 1.5 mm creating a 60° light beam that covered the full diameter (1.5 cm) of the craniotomy (at 40-62 mW power). **D**. Blue light stimulation (451 nm) at 10Hz with a 50% duty cycle (50 ms pulse width) with 30-sec long stimulation on/off block pattern. **E**. BOLD signal modulation and power spectrum of an example voxel in V1 along with the voxel modulation peak at 0.016 Hz or 1 cycle/60 sec. **F**. In-volume activation map of monkey M1 with the yellow brackets indicating the chamber region. **G**. Pial surface reconstruction of each individual monkey showing the model-free activation map based on the coherence between each voxel BOLD signal modulation and the stimulation paradigm (coherence threshold > 0.3). The extent of V1 cortical activation (50 -120 mm^2^) is summarised in supplementary **Table 1**. **H**. Summary of the local response clustered area across three of the macaque monkeys (VL, DP and AL) shown on the inflated surface of the D99 macaque template. **I**. Post-mortem brain of monkey FL showing the native eyfp fluorescence after surface light stimulation with blue light on the opercular region of V1. **J**. Plot of the mean and +/- std time course of each voxel as percent signal change obtained from the local clustered region of monkey VL for both blue (451 nm; blue trace) and red-light (620 nm; red trace). Note the wavelength dependence of the BOLD modulation **K**. Electrophysiological based optogenetic modulation of the transfected opercular region in V1 of monkey VL. Top left panel shows the schematic approach for recordings during opto stimulation. Multiunit activity during 15 sec (on and off) optogenetic stimulation delivered at 10 Hz. Individual plots show the multiunit activity across channels located at different depths perpendicular to the cortical sheet.

**Table 1.**

|  | *Blue light* |  | *Red light* |  | *Movie* |  |
| --- | --- | --- | --- | --- | --- | --- |
| *Monkey* | Activation<br>cluster<br>mean<br>coherence | Activation<br>cluster<br>sigma<br>coherence | Activation<br>cluster<br>mean<br>coherence | Activation<br>cluster<br>sigma<br>coherence | Activation<br>cluster<br>mean<br>coherence | Activation<br>cluster<br>sigma<br>coherence |
| *Velma* | 0.4 | 0.2 | 0.14 | 0.08 | 0.62 | 0.19 |
| *Daphne* | 0.55 | 0.28 | - | - | 0.33 | 0.18 |
| *Alma* | 0.45 | 0.16 | 0.21 | 0.09 | 0.31 | 0.09 |
|  | *Blue light* |  | *Red light* |  | *Movie* |  |
| *Monkey* | Activation<br>cluster<br>mean<br>phase | Activation<br>cluster<br>sigma<br>phase | Activation<br>cluster<br>mean<br>phase | Activation<br>cluster<br>sigma<br>phase | Activation<br>cluster<br>mean<br>phase | Activation<br>cluster<br>sigma<br>phase |
| *Velma* | 196.78 | 30.62 | 151.53 | 44.2 | 172.33 | 35.62 |
| *Daphne* | 150.22 | 23.15 | - | - | 147.95 | 24.46 |
| *Alma* | 103.23 | 26.81 | 131.2 | 35.37 | 214.91 | 39.83 |

| **Monkey** | **Total**<br>**number**<br>**active**<br>**voxels** | **Total**<br>**number of**<br>**V1 voxels** | **V1**<br>**volume**<br>**(mm3)** | **Percent**<br>**activation**<br>**of V1**<br>**volume** | **Activation**<br>**volume**<br>**(mm3)** | **voxel size**<br>**mm3** |
| --- | --- | --- | --- | --- | --- | --- |
| *Velma* | 1840 | 72199 | 1949.37 | 2.55% | 49.68 | 0.3 |
| *Daphne* | 2816 | 71383 | 1927.34 | 3.94% | 76.03 | 0.3 |
| *Alma* | 3709 | 83422 | 2252.4 | 4.45% | 100.43 | 0.3 |

| **Monkey** | **Total**<br>**number**<br>**active**<br>**surface**<br>**voxels** | **Total**<br>**number of**<br>**surface**<br>**V1 voxels** | **V1 area**<br>**(mm2)** | **Percent**<br>**activation**<br>**of V1 area** | **Activation**<br>**area**<br>**(mm2)** | **voxel size**<br>**mm2** |
| --- | --- | --- | --- | --- | --- | --- |
| *Velma* | 535 | 11493 | 1034.27 | 4.65% | 48.15 | 0.3 x 0.3 |
| *Daphne* | 808 | 11401 | 1026.09 | 7.08% | 72.72 | 0.3 x 0.3 |
| *Alma* | 1354 | 12585 | 1132.65 | 10.75% | 121.86 | 0.3 x 0.3 |

| **Monkey** | **Activation**<br>**cluster**<br>**mean**<br>**%CNR** | **Activation**<br>**cluster**<br>**sigma**<br>**%CNR** | **Activation**<br>**cluster**<br>**mean**<br>**%CNR** | **Activation**<br>**cluster**<br>**sigma**<br>**%CNR** | **Activation**<br>**cluster**<br>**mean**<br>**%CNR** | **Activation**<br>**cluster**<br>**sigma**<br>**%CNR** |
| --- | --- | --- | --- | --- | --- | --- |
| *Velma* | 0.7 | 0.1 | 0.4 | 0.2 | 2.0 | 0.12 |
| *Daphne* | 0.76 | 0.14 | - | - | 1.6 | 0.4 |
| *Alma* | 0.94 | 0.2 | 0.05 | 0.1 | - | - |

The effectiveness of the elicited optogenetic stimulation was then first assessed during opto-fMRI experiments using a 4.7 T vertical MRI scanner acquiring images at high-resolution (1.2 mm isotropic voxel size). Blue light stimulation parameters consisted of 451 nm wavelength set at 10Hz with 50% duty cycle and a power range of 40-62 mW (**Supp. Fig. 2A**). Light was delivered in alternations of 30-sec long on and off blocks, while the monkeys were at rest (**Fig. 1D**). Examination of the BOLD signal time course in a voxel located at the V1 chamber position revealed its systematic modulation during periods of optogenetic stimulation (see left panel on **Fig. 1E**). In order to quantify the BOLD modulation in response to optogenetic stimulation, we used voxel-based coherence analysis as the analytical method of choice, since the hemodynamic response function (HRF) of opto-fMRI in macaques has not been formally characterized. To this end, the strength of the BOLD response was assessed by first calculating the frequency-resolved power-spectrum and then determining the coherence between the BOLD peak frequency response (cycles/volume) and the stimulus repetition rate (0.016 Hz = 1/60 second) (see right panel in **Fig. 1E**).

From the opto-fMRI experiments, we observed significant BOLD modulation (cluster size > 10 voxels, coherence > 0.35, n runs = 4, see coherence map on raw EPI volume and without threshold in **Supp. Fig. 3B**) in response to blue-light stimulation of the opercular area of V1 (**Fig. 1F**, for threshold coherence map overlaid on T1 anatomy). Comparison with visual stimulation (**Supp. Fig. 3A**) revealed that opercular V1 was also active during natural vision and that visual stimulation resulted in ∼0.5% stronger BOLD modulation compared to optogenetic stimulation under these experimental conditions. To better visualize the activation pattern elicited in V1 by optogenetic stimulation, we overlaid the BOLD activation map on a 3D surface reconstruction of the occipital lobe (**Fig. 1G**). Across the three imaged monkeys, the local activation mapping revealed an activation focus over the dorsal opercular V1, with a size of 50 - 120 mm^2^ corresponding to about 5 - 10% of the V1 surface area (see **Table 1**). Interestingly, in all three monkeys, weaker activation could also be observed in the more ventral part of V1, possibly reflecting the local dorso-ventral connectivity (Casagrande and Kaas, 1994). The spatial proximity of the activated V1 zones across the three monkeys (VL, DP and AL) is shown in the standardized D99 macaque brain template (Reveley et al., 2017) in **Fig. 1H** and compared to the *ex-vivo* fluorescence pattern from an additional monkey (FL) in which a similar injection approach was performed (**Fig. 1I**). Thus, these initial results demonstrated the effectiveness of the optogenetic approach in driving the local V1 BOLD response using fMRI. Our next aim was then to establish the extent to which the observed BOLD response pattern was wavelength-specific and whether optogenetic stimulation could also drive local spiking activity.

### V1 BOLD and neural responses to 451 nm but not to 626 nm stimulation

An important confound to consider in optogenetics is the potential for heat buildup in the brain tissue due to light absorption (Stujenske et al., 2015). In particular with opto-fMRI, heating induced negative BOLD signal modulations have been observed in rodent animal models (Christie et al., 2013; Desai et al., 2010). While we observed only positive BOLD signal modulation during optogenetic stimulation, we attempted to address the potential effects of heat by carrying out additional control experiments.

With the first control, we aimed to determine if the local BOLD response was specific to blue-light stimulation (451 nm), or if the response could also be evoked by red light (626 nm) stimulation. We assumed that the BOLD responses should only be driven by blue-light to which ChR2 is sensitive (Boyden et al., 2005). In contrast, stimulation with red light should result in no response given that ChR2 is not sensitive to 626 nm wavelength light stimulation. To address the potential confound of wavelength-specific effects, we carried out experiments in the same ChR2-transfected animals. During experiments we alternated between blue and red light stimulation runs keeping the stimulation power constant at 50 mW and pulsed at 10 Hz, 50% duty cycle for both wavelengths (**Supp. Fig. 2B**). In monkey VL (session M31), we found again a local positive BOLD response in V1 to blue light (451 nm, 50 mW at 10Hz; cluster size > 10 voxels, coherence > 0.35, n runs = 4; **Fig. 1J,** blue line; see coherence map without threshold in **Supp. Fig. 4A and C**). During the same session, but on alternating runs, we also stimulated with red-light. Here, we found no activation (626 nm, 50 mW at 10Hz; cluster size > 10 voxels, coherence > 0.35, n runs = 4; **Fig. 1J,** red line; see coherence map without threshold in **Supp. Fig. 4B**) at the same local V1 site previously activated by blue-light. On average across experimental runs, the optogenetic target region in V1 displayed lower contrast-to-noise ratios (CNR) for red-light (mean CNR %, 0.4 ± std 0.2) as compared to blue light (mean CNR %, 0.7 ± std 0.2). Similarly, blue light stimulation resulted in a significant coherence activation cluster (n runs = 6, coherence of 0.4, ± std 0.2), while red light resulted in non-significant modulation (n runs = 6, coherence of 0.14, ± std 0.08). To assess the reliability of this effect, we carried out additional opto-fMRI experiments in monkey AL and found significant local BOLD modulation (see map without threshold and with threshold in **Supp. Fig. 4D;** cluster size > 10 voxels, coherence > 0.35, n runs = 4) only for blue-light (451 nm, 50 mW at 10Hz) but not for red-light (626 nm, 50 mW at 10Hz). Similarly, only blue-light stimulation resulted in significant BOLD modulation (n runs = 4, mean coherence 0.45, ± std 0.21, mean CNR % 0.94 ± std 0.2) while red-light resulted in no response modulation (n runs = 4, mean coherence 0.21, ± std 0.09, mean CNR % 0.05 ± std 0.1).

To further investigate the neurophysiological basis of the observed optogenetically induced BOLD fMRI response in V1, we inserted multi-contact electrode arrays into V1 and recorded the spiking activity of cortical neurons during optical stimulation of the transfected brain tissue. Optical stimulation was delivered epidurally as similarly performed during fMRI experiments (**Fig. 1K**). To represent neuronal spiking activity, we extracted one multiunit for each channel. In monkey VL, ten sites were sampled throughout five experimental sessions. In total, 104 units were extracted, 85 units showed significant increase in firing rate in response to a one-second continuous pulse stimulation with blue light (451 nm, p = 0.05, Wilcoxon signed rank test). The units extracted covered the whole depth of the cortex showing reliable neural activation across the V1 target zone. The response latencies of the multi-unit activity (MUA, see **Methods**) were short (< 10 ms) indicating a direct activation by blue light; characteristic of ChR2 (Boyden et al., 2005). Additionally, we observed increases in spiking in the V1 target site only in response to stimulation with blue (451 nm), but not red (626 nm) light. In summary, with our initial experiments we were able to determine that the local BOLD response underlying the injection sites reflected changes in neural activity specific to blue light and that heat-related effects were rather unlikely.

### V1 neural activity closely matches layer specific immunohistological expression

To further examine the effects of optogenetic modulation on the V1 cortical microcircuit, we inserted layer resolving multi-contact electrode arrays into the V1 of monkey FL and recorded the spiking activity of cortical neurons during optogenetic stimulation of the transfected brain tissue. The laminar coverage of the probes was determined based on the current source density (CSD) profile of the local field potential (LFP) and the MUA across channels in response to a visual stimulus. From these measures we identified the geniculate recipient layer 4C as the electrode contacts that carried the earliest current sink (**Supp. Fig. 6A**) and shortest MUA response latency (< 50 ms, **Supp. Fig. 6B**) (Maier et al., 2010; Schroeder et al., 1998).

For optogenetic stimulation, the laminar electrode was equipped with an embedded optical fibre allowing intracortical light delivery at 350-650 µm, as estimated from the relative cortical depth in monkey FL. Across 19 sessions, 354 out of 380 multi-units showed significant increase in firing rates (p < 0.05, Wilcoxon signed rank) in response to a continuous 300 ms stimulation pulse with blue light (473 nm) delivered via laser (see **Methods**). As expected, red-light (594 nm) did not affect neural activity (**Supp. Fig. 8**). Across experimental sessions, we sampled from locations up to 2 mm around the injection site. The averaged laminar profile of the sustained (100 - 300ms) MUA response showed significant increases in firing rates across sessions (19/20 contacts p < 0.001, and one contact p = 0.05, Wilcoxon signed rank, right tailed test) with activation peaks in putative layers 4B followed by layers 5 and 6 (**Fig. 2A,** blue line). Histological analyses of the V1 transfected region (**Supp. Fig. 1E)** confirmed an increase in percentage of eYFP positive cell expression in layers 4B, 5 and 6 with sparser expression in layers 2/3 (**Fig. 2B**) resulting in a significant correlation between the electrophysiological response and histological expression profiles (**Fig. 2B**, Spearman’s ⍴ = 0.5273, p = 0.026).

**Figure 2.**
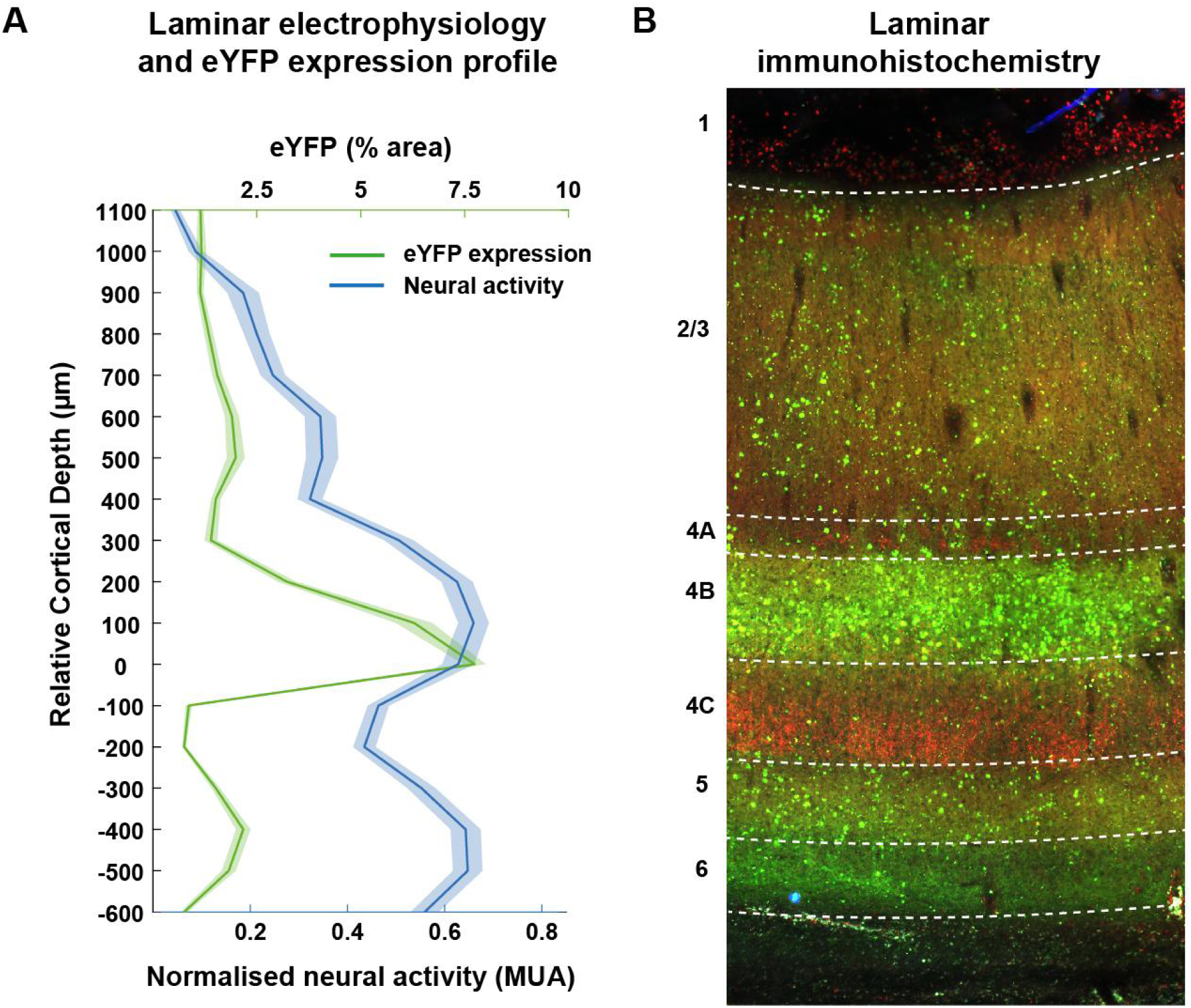
V1 neural activity closely matches immunohistochemistry expression profile across cortical layers. **A**. Sustained (100-300ms) laminar activation pattern (blue) for continuous (300 ms) optogenetic stimulation and area percentage expressing eYFP (green) as a function of relative cortical depth (aligned to layer 4C). Alignment was calculated based on the earliest response to a visual stimulus. **B**. Laminar profile of V1 near the injection site. eYFP expression of the optogenetic construct can be seen in green, layer 4C (particularly 4C beta) was co-stained using an anti-vGlut2 antibody using standard immunohistochemistry (red)..

Taken together, our results with fMRI, electrophysiology and histology demonstrate a robust activation of the local V1 microcircuit, including strongest response modulation in layer 4B that connects V1 to extrastriate cortical areas (Casagrande and Kaas, 1994). Somewhat surprisingly, however, our initial fMRI assessment showed very little remote activation beyond V1. As frequency dependent effects have been reported in studies using electrical stimulation in combination with neuroimaging (es-fMRI (Kampe et al., 2000; Logothetis et al., 2010; Murris et al., 2020; Van Camp et al., 2006), we reasoned that optogenetic stimulation frequency could be a mediating factor in driving activation into higher-level regions

### Increasing V1 stimulation frequency to 40 Hz drives extrastriate BOLD activity

We tested the effect of applying different optogenetic stimulation frequencies (5 Hz, 10 Hz, 40 Hz, 50% duty cycles) suited to activate ChR2 (Boyden et al., 2005) on the V1 laminar electrophysiological response level, in addition to determining frequency dependent BOLD response modulation. As predicted, we found that neural activity followed the optogenetic stimulation frequency closely (**Fig. 3A**). To evaluate the response pattern further, we then calculated an averaged laminar activation profile for each frequency as previously done for continuous stimulation. For both 5Hz and 10Hz, the activation increases were small and uniform, and the laminar activation pattern was not different from continuous stimulation (**Supp. Fig. 7**). However, stimulation delivered at 40 Hz caused an overall increase in activation and especially in the more superficial layers, which was not observed with continuous or lower frequency stimulation (**Fig. 3B**). As superficial layers are known to send their projections to higher order areas of visual association cortex (Casagrande and Kaas, 1994), we took advantage of fMRI to map whether 40 Hz stimulation was more efficient in driving BOLD activation remotely.

**Figure 3.**
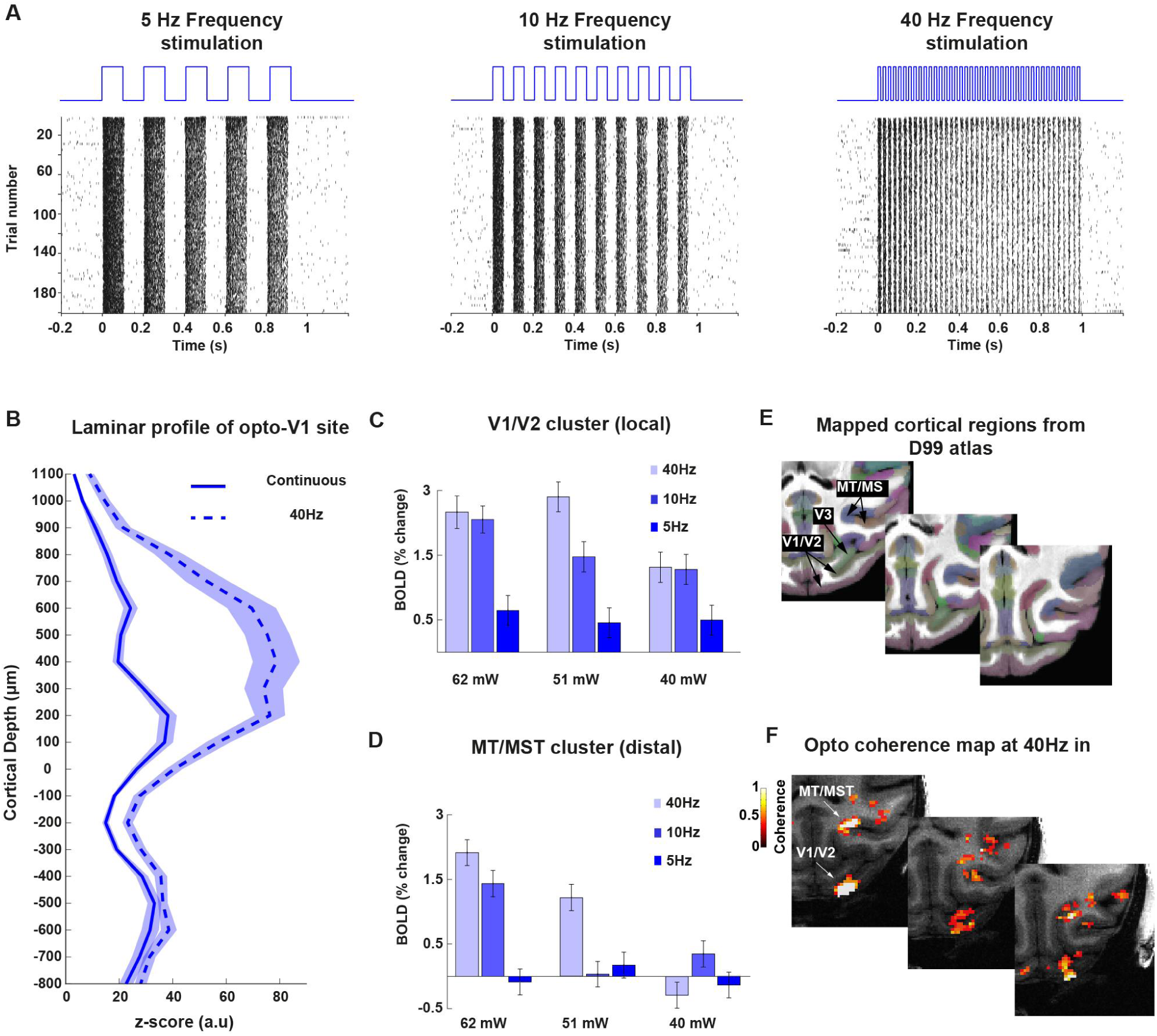
Increases in extrastriate BOLD activity after increased frequency stimulation of opercular V1. **A**. Example raster plots of neural activation shows reliable temporal modulation of spiking activity from optogenetic stimulation rate (5Hz, left panel; 10 Hz, middle panel, and 40Hz right panel). **B**. Laminar profile of the local V1 neural activation during continuous (solid line) and 40 Hz stimulation (dashed line). The z-score reflects the neural activation strength as a function of cortical depth centered at the granular level (layer IV or 0 distance in mm). The pattern shows increased spiking modulation (mean z-score +/- std) in supragranular layers of the transfected region in V1 as compared to continuous stimulation (blue line). **C**. Average percent BOLD signal change of the V1/V2 local cluster region tested at three different power levels (40 mW, 51 mW and 62 mW) and three different frequency levels (5Hz, 10Hz, and 40Hz). **D**. Average percent signal change of the MT/MST distal cluster region as similarly shown in **C.** Note the strongest modulation occurs at maximum power levels and at 40 Hz frequency. **E**. Cortical regions form the D99 atlas (top panel) mapped onto the warped template of monkey VL. **F**. Maps of monkey VL showing significantly active regions which are labeled as follows: visual areas V1, V2 and V3 and motion-sensitive regions MT/MST. Similar maps on monkey DP are showing in **Supp.Fig. 10. A**.

For these opto-fMRI measurements, the global stimulation design was kept constant at 30 seconds (ON and OFF), while the pulse rate was set at 50% duty cycle for all the frequencies (5 Hz, 10 Hz, and 40 Hz) and different power levels (40 mW, 51 mW and 62 mW) were tested. To identify active regions driven by the optogenetic stimulation we overlaid the D99 atlas (Reveley et al., 2017) onto the warped template of our in-session anatomical scan (**Fig. 3C**). Focusing on early striate and extrastriate cortex, we observed activation in areas V1/V2 and MT/MST under 40 Hz stimulation (**Fig. 3D** and **Supp. Fig. 7**). Closer examination of the response magnitude revealed effects for both stimulation amplitude as well as frequency (**Fig.3 E and F**). To drive BOLD activity in area MT more than 40 mW was necessary during epidural stimulation (**Fig. 3C** and **D**). Importantly, at the strongest stimulation power of 62 mW and 40 Hz stimulation frequency, the BOLD response effect was more efficient than at lower stimulation frequencies. Our finding of increased opto-fMRI activation with higher stimulation frequencies is consistent with previous studies using electrical microstimulation in combination with fMRI (Kampe et al., 2000; Logothetis et al., 2010; Murris et al., 2020; Van Camp et al., 2006).

### Predominant dorsal stream BOLD activity after optogenetic stimulation of V1

Having established the effectiveness of 40 Hz stimulation for driving local and remote activation, we wondered in which areas of visual association cortex activation could be measured with fMRI and how robust such activation would be in our different monkeys. **Figure 4A** shows the overall BOLD activation pattern in monkey FL in a cortical flat map for better visualisation. Optogenetic stimulation was effective in driving activation in several cortical areas of the visual system, including V1, V2, V3, MT/MST and FEF (**Fig. 4B).** Additional activation could also be observed subcortically in the lateral geniculate nucleus (LGN) (**Supp. Fig. 10).** Despite slight variations with respect to the specific dorsal part of V1 being optogenetically transfected and stimulated, qualitatively very similar results with activation of extrastriate cortex were observed across the three monkeys (see **Supp. Fig. 9** for flat maps on individual monkeys). This similarity in the cortical activation pattern is highlighted in the averaged activation map across the three investigated monkeys with predominant activation of areas belonging to the cortical dorsal stream and parietal cortex (**Fig. 4C and E).** Interestingly, despite the previously noted activation in ventral V1/V2, only limited activation could be observed in ventral stream regions beyond V1/V2 and the temporal lobe. These results from opto-fMRI appear qualitatively very similar to previous tracer-based characterization of V1 connectivity REF. As this optogenetically induced response map with predominant dorsal stream activation only partially reflects the rich cortical activation pattern normally seen during free-viewing conditions (see **Supp. Fig. 3B**), we wondered whether the recruited network of brain areas might be sufficient to induce an artificial visual percept.

**Figure 4.**
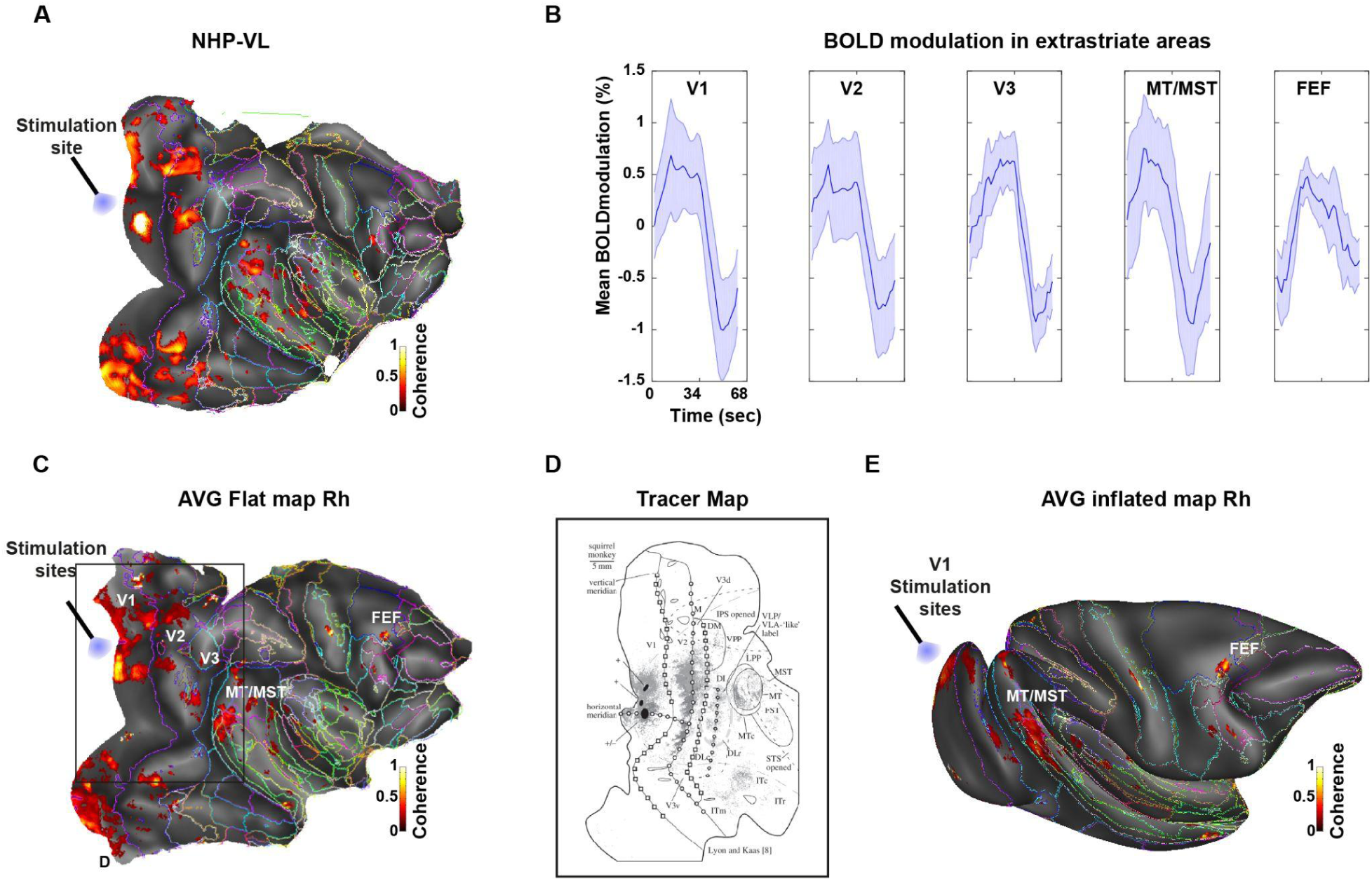
V1 optogenetic stimulation drives BOLD activation in higher cortical areas. **A**. Cortical activation map of monkey VL in response to optogenetic stimulation of opercular V1. **B**. Average single-cycle BOLD modulation across activated cortical regions showing extrastriate BOLD activity in V1, V2, V3 and motion complex regions MT/MST in example monkey VL with additional activity seen in area FEF. For additional maps in monkey DP and AL see **Supp.Fig 9**. **C** and **E**. shows flat and inflated surfaces, respectively, for the averaged activation map across all three monkeys (coherence > 0.3, mean 0.46 > 0.2 +/- std). The map shows cortical regions driven by optogenetic stimulation of opercular V1. LED fiber points to the stimulation sites in the opercular region, with a size of 50 -120 mm^2^ corresponding to about 5 - 10% of V1. Activation outside V1 includes motion-sensitive regions MT/MST and area FEF in the frontal lobe. **D.** V1 tract-tracing map (Lyon and Kaas, 2002) showing a terminal labelling pattern in new world monkeys that is very similar to the activation map obtained with our opto-fMRI in macaques (panel C).

### V1 optogenetic stimulation induces a visual “phosphene”

To probe the behavioral effects of V1 optogenetic stimulation, we trained monkey FL to perform visual perceptual reports using a two-alternative forced-choice task (2AFC, **Fig. 5A**). The monkey had to initially fixate on a dot at the center of a computer monitor (500-1250 ms). After successful fixation, one of three equally probable conditions (optogenetic stimulation with a continuous light pulse, visual stimulation at one of five possible luminance levels, catch trials without stimulation) followed. Before starting with the series of optogenetic experiments, the monkey had been trained to report its percept irrespective of stimulus position by making a saccade to the left of two targets, indicating the presence of a stimulus. During catch trials the animal was required to make a saccade to the target on the right instead of the left, effectively reporting the absence of a stimulus. In both cases, the animal received an equal amount of liquid reward. Once the monkey reliably reported its visual percept with a hit rate of > 80% for both high-contrast visual stimulation and catch trial conditions, optogenetic stimulation of V1 was introduced. A critical component of the experimental procedure was to first electrophysiologically map the RF of optogenetically targeted neurons, before proceeding with the behavioural testing at this opto-RF location. In an example session, the monkey’s overall performance during optogenetic stimulation was at 86% and therefore as high as during visual stimulation and catch trials (**Fig. 5A**). Comparing the performance during the opto trials with those during different visual luminance conditions demonstrated that the monkey had, on average across experiments, a sensitivity (d’, see **Methods**) of about 1.3 for optogenetic stimulation, matching the sensitivity to a 10% visual stimulus luminance increase (**Fig. 5C**). Thus, the monkey’s report from optogenetic stimulation closely resembled his reports when a low contrast visual stimulus was present.

**Figure 5.**
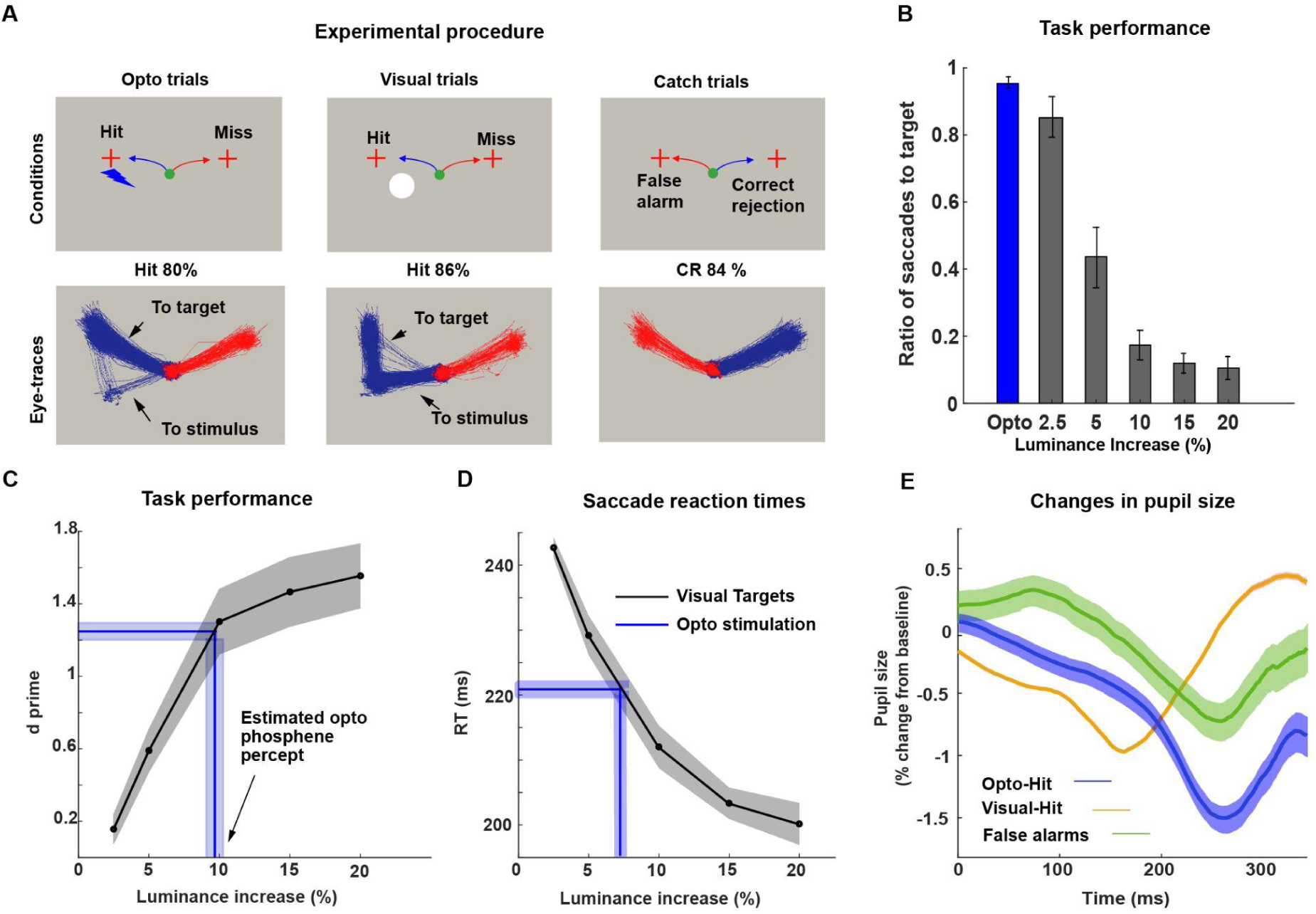
Induction of a visual phosphene from optogenetic stimulation of primate V1. **A**. Schematic representation of the 3 experimental conditions (top panel) and associated eye movement trajectories in response to each stimulation condition (bottom panel). Note that optogenetic stimulation typically resulted in direct saccades to the response target, whereas saccadic reactions to visual stimulation were typically first directed at the visual stimulus, before continuing to the response target. **B**. Proportion of saccades to target vs to stimulus locations across the different experimental conditions. The majority of eye movements in response to optogenetic stimulation were directed to the target. The number of saccades to target decreased with increasing luminance levels in the visual condition. **C** and **D**. Comparison of visual sensitivity **C** and saccadic reaction times **D** between the visual and optogenetic stimulation conditions. The blue line indicates the mean and SEM visual sensitivity to optogenetic stimulation. Projection of optogenetic to visual sensitivity and reaction time curves indicates that the optogenetically elicited phosphene likely corresponds to the percept evoked by a 5 - 10% luminance (increase over background) visual stimulus. **E**. Comparison pupil size changes following visual (orange line) vs optogenetic (blue line) stimulation and catch trials (green line). Shading indicates standard error of means.

In order to establish whether a similar effect was also observable under implicit, physiological measures, we examined closely the monkey’s eye movement response pattern (**Fig. 5A** and **5B**). The testing regimen allowed the monkey to either make a saccade to the stimulus first and then to the response target, or straight to the target for both visual and opto conditions. In the visual condition, the macaque’s saccade trajectory differed according to the luminance of the visual stimulus; under low contrast conditions the monkey typically looked directly at the response target, whereas under high contrast conditions the monkey first looked at the visual stimulus before making an additional saccade to the response target. In the opto condition, the majority of the monkey’s saccades were aimed directly towards the response target, highlighting again the close similarity between performance during opto and low luminance contrast visual trials. Similarly, reaction times of saccades (**Fig. 5D**) after visual stimulus onset decreased with increasing luminance and the reaction times of saccades after optogenetic stimulation were similar to low contrast visual stimulation. In addition to these observations on eye movements, we also unexpectedly noted a pupillary constriction in response to optogenetic stimulation (**Fig. 5E**) that occurred with a ∼100 ms longer latency compared to visually induced pupillary constriction.

Taken together, the data from one monkey demonstrate that V1 optogenetic stimulation can induce an artificial visual percept that appears to resemble the percept of a low contrast stimulus.

## Discussion

Despite the increasing importance of establishing optogenetic methods for investigations in non-human primates, the ability to initiate behaviour and to cause changes in perception in this species remains a challenge. By performing a multi-modal assessment of our V1 optogenetic stimulation approach in macaque monkeys, we confirm the effectiveness of optogenetic stimulation for driving the V1 laminar circuitry and higher-order visual cortical areas, as well as for inducing artificial visual percepts (‘phosphenes’). In our discussion, we highlight the benefits and limitations of using optogenetics for mapping the V1 microcircuitry and the large-scale cortical networks, before considering how this method can be optimally used to induce meaningful behaviour. We compare our findings with existing studies in which macaque V1 has been stimulated either with optogenetics or with electrical microstimulation.

### How does optogenetic stimulation interface with the V1 microcircuit?

One of the key elements of optogenetics is that it enables the interference with specific compartments of the neural microcircuitry to an extent that is presently not achievable with more traditional electrical microstimulation techniques. Whereas electrical stimulation is thought to primarily affect bypassing axonal projections (Histed et al., 2009), the development of opsins, promoters, viral vectors, as well as wavelength-specific light delivery systems can be harnessed for targeting specific circuit elements. While the cortical microcircuit organization follows many common principles across regional boundaries, one of the challenges for optogenetic methods in NHPs is drawing common conclusions about its efficiency across studies targeting different brain regions (Tremblay et al., 2020). At the level of primate V1 few studies have started to delineate more systematically the laminar expression pattern of optogenetic constructs and how optogenetics affects functional architecture. Following injections into primate V1 among other areas, Gerits et al. reported serotype specific expression patterns (Gerits et al., 2015). Their tests did however not include the AAV9-hSyn construct applied in our study, yet at the same time none of their tested constructs under CamKII and CMV promoter control exhibited such a strong laminar expression specificity as in our study. A V1 laminar expression profile similar to the one in our preparation was reported in an earlier study in marmosets and macaques, in which the hSyn promoter was examined (Watakabe et al., 2015). Again a much less specific laminar distribution was observed for the CamKII promoter. The authors further reported that the neurons with strong expression under the hSyn promoter had the appearance of pyramidal neurons. Consistent with this laminar specific profile for hSyn, Jazayeri et al., who injected AAV1-hSyn-ChR2(H134R)-mCherry, also reported a very similar laminar expression pattern to that in our study, with strongest expression in layers 4B, 5 and 6. As neurons in these layers contribute to V1 output connectivity with other areas, including layer 4B to motion sensitive area MT, layer 5 to pulvinar and superior colliculus and layer 6 to LGN (Casagrande and Kaas, 1994), the fMRI activation that we found in these areas corresponds well to the observed expression pattern. Our study, is consistent with earlier reports, which also found an expression gap in layer 4C, where parvo- and magnocellular lateral geniculate nucleus (LGN) axonal projections terminate, which we visualized by means of vGlut2 staining as a selective marker for this geniculate input into V1 (Garcia-Marin et al., 2013). From this review of the current literature and our present findings, a picture emerges that emphasizes the greater laminar specificity with hSyn compared to CamKII promoters, at least at the level of V1, opening the door to use targeted light delivery to these layers to investigate their connectivity and function for vision. Along those lines, two recent studies have successfully implemented microscopy and artificial dura preparations to image the *in-vivo* fluorescence of their injected lentivirus-based optogenetic constructs along the cortical surface and have started to delineate functional implications: In addition to demonstrating highly confined topographic expression, Nassi et al showed how the interaction between local excitatory neural networks shapes neuronal normalization in electrophysiological measures (Nassi et al., 2015). Chernov et al managed to image the columnar specificity of optogenetic construct expression and how stimulation of one column leads to the selective activation of more distant columns with related function (Chernov et al., 2018). A limitation of our study is that we did not determine the functional specificity of our optogenetically identified neurons. Addressing functional specificity would be important to clarify our knowledge on the laminar distribution of visual computations of V1 neurons that could not draw yet on optogenetic labeling technologies. We expect that neurons in layer 4B, where we observed strongest expression in our preparation, to display strong orientation and direction of motion tuning (Gur et al., 2005; Movshon and Newsome, 1996). Knowledge of the neurons that can be targeted with optogenetic stimulation is also important to establish what kind of V1 processing can be harnessed effectively to modulate or shape a visual percept.

### Mapping brain connectivity using opto-fMRI

An important study by Tolias et al reported for the first time the successful implementation of electrical microstimulation in combination with fMRI (‘es-fMRI’) as a method to map effective brain connectivity (Tolias et al., 2005). From stimulating V1, the authors reported BOLD activation in the extrastriate cortex which closely resembles our findings with optogenetic stimulation (**Fig. 4**), including activation of V2/V3 and MT/MST. Consistent with our findings is also the limited activation of area V4 and of further ventral stream regions as similarly observed by the Tolias study using es-fMRI. In addition to this parieto-occipital activation pattern, we also observed activation in the frontal eye field (FEF). We attribute the lack of FEF activation in the Tolias study to the selection of RF coils or to the use of anaesthesia, which often tends to suppress activation especially in high level association cortex. As V1 connects with all three clusters, V2/V3, MT and FEF (Casagrande and Kaas, 1994; Perkel et al., 1986), it is conceivable that our activation pattern primarily reflects monosynaptic connectivity predominantly within the cortical dorsal stream. Such an interpretation is consistent with the findings of the Logothetis lab that applied electrical stimulation to LGN and measured associated cortical responses and concluded that es-fMRI affects inhibitory circuits which prevent multi-synaptic propagation of action potentials (Logothetis et al., 2010). These findings, however, contrast with many other reports in which polysynaptic effective connectivity could be elicited with es-fMRI (Ekstrom et al., 2008; Moeller et al., 2017).

On a more technical note, a limitation of our fMRI results is that we can not entirely rule out heating effect contributions (Stujenske et al., 2015). While our comparison of the effects of blue vs red light stimulation on the BOLD response and electrophysiological signals make heating effects appear unlikely, we can not entirely exclude such heating contributions to our fMRI results. fMRI measurements under blue light stimulation in non-transfected cortex would have been a better control for heating (Stujenske et al., 2015). It is also important to bear in mind that our study is only the second to report successful use of opto-fMRI in NHP, which contrasts with the routine use of optogenetics in non-primate species. The study by Gerits et al successfully used optogenetic stimulation in FEF to map the large-scale effective connectivity from area FEF and successfully influenced saccadic eye-movements. In contrast, a study with a seemingly identical approach for FEF stimulation, found no optogenetically induced fMRI activation, though it was possible to drive neuronal responses and influence saccadic behaviour. Furthermore, optogenetic stimulation appeared to reduce activation elicited by es-fMRI. Why might these two studies have come to very different fMRI results despite their seemingly very similar experimental approach? Close examination of the applied experimental parameters reveals that Gerits et al used 40 Hz optogenetic stimulation, whereas Ohayon et al applied brief continuous stimulation pulses for optogenetic stimulation and applied 300 Hz stimulation only under es-fMRI conditions. Our findings here indicate that pulsed 40 Hz stimulation is more effective in driving electrophysiological activity in V1 superficial layers and for propagating activation to remote brain areas (**Fig. 3**). The benefit for high frequency stimulation is well known from electrical stimulation studies (Kampe et al., 2000; Logothetis et al., 2010; Murris et al., 2020; Van Camp et al., 2006) and might help to explain the difference in mapping effective connectivity between the two previous NHP opto-fMRI studies.

### Using optogenetics in V1 to induce a visual phosphene

In stark contrast to the many studies employing electrical stimulation techniques with great success, only very limited evidence exists in NHPs that demonstrates the capacity to affect behaviour from optogenetic stimulation of cortex. This is exemplified by the small number of reports in which vision guided behaviour could be successfully manipulated using optogenetic methods, despite the great anatomical accessibility of V1 and the clear-cut testable predictions for behavioural consequences of its manipulation. The pioneering study by Jazaheri et al using ChR2 demonstrated how spontaneous eye movements tended to be directed to the receptive field of optogenetically identified neurons. While this finding corroborated earlier observations obtained with electrical stimulation (Tehovnik et al., 2003), a limitation was that it remained unclear what the monkey perceived from such stimulation. While a recent study did not report whether eye movements or a percept could be elicited from optogenetic stimulation of V1, the authors demonstrated how the stimulation could improve the detectability of weak visual stimuli, provided there was a match in the stimulus properties and the tuning for this property of the optogenetically targeted neurons (Andrei et al., 2019). Our laminar stimulation approach might have contributed to our success in eliciting a visual phosphene. Finally, a study by De et al recently showed how optogenetic V1 inactivation disrupted eye movements and reduced stimulus visibility by more than 50% during a perceptual choice task. Here our results in one monkey (**Fig. 5**) fill in an important gap in the literature; stimulating V1 optogenetically induced a visual percept (‘phosphene’) which the monkey reliably reported with a hit rate over 80%. Interestingly, in addition to its effect on explicit perceptual reports, optogenetic stimulation also induced pupillary constriction, which we interpret as a cognitive effect due to its increased latency by about 100ms, and attribute to the recruitment of cortical networks implicated in the control of pupillary responses (Peinkhofer et al., 2019). Gaining access to such implicit performance measurements might help in future studies aimed at identifying behavioural consequences from optogenetic stimulation. For practical reasons, we had initially tried to perform behavioural tests of optogenetic stimulation without additional electrophysiology, yet without much success. For one, we felt it was easy to induce behavioural biases unless meaningful controls such as catch trials were included. In addition, it proved difficult for us to be certain about the spatial location of the expected optogenetic effect. This changed after we had mapped the opto-RF and implemented 2AFC methods, giving us greater certainty over the measured effects. Nevertheless, our approach contains limitations, including the reward schedule, the timing of events which could have contributed to our effects. In an effort to better understand the perceptual quality of the evoked phosphene, we related the monkey’s performance during optogenetic stimulation to his response pattern under visual stimulation at varied luminance contrast levels. The performance measurements along with the eye movement patterns indicate that the monkey most likely perceived a phosphene that resembles a 10% luminance contrast visual stimulus. This finding is very similar to observations made with electrical stimulation under comparable test conditions (Schiller et al., 2011). Naturally the question then arises how such a phosphene could be strengthened and how it could be turned into something more useful for visual function. Based on our assessment of stimulation frequency and amplitude effects on the recruitment of activation in higher-order association areas, we believe it might be preferable to apply 40 Hz stimulation also for behavioural testing. Another clear prediction from our results of optogenetic expression in layer 4B which projects to motion-sensitive area MT, where we reliably observed activation in fMRI, is the possibility to induce a percept of motion, as has been also recently demonstrated with electrical stimulation and current steering methodology (Beauchamp et al., 2020b).

Taken together, our results highlight some of the current challenges, but also promises, that optogenetic applications for the primate brain face. Using advanced technologies that increasingly also target the topographic domain of the sensory-motor cortex, it should be possible to recruit neural circuits more efficiently and elicit meaningful behaviour.

## Materials and Methods

### EXPERIMENTAL DESIGN

#### Subjects

Four female rhesus monkeys (*Macaca mulatta*) were used to obtain all opto-fMRI data; (VL, six years of age, weighing 7 kg; DP, six years of age, weighing 9 kg; AL, five years of age, weighing 9 kg; FL, 4 years of age, weighting 6 kg). Surgical and anesthesia procedures, postoperative care, and implant methods were described in detail in a previous manuscript (Ortiz-Rios et al., 2018). The UK Home Office approved all procedures, and procedures comply with the Animal Scientific Procedures Act (1986) on the care and use of animals in research and the European Directive on the protection of animals used in research (2010/63/EU).

#### Viral injections

We used the Adeno-Associated Virus (AAV) serotypes 5 and 9 as the viral vector for gene delivery for all our constructs injections (n = 4 monkeys, all injected in the right hemisphere, see **Supp. Fig. 1A** for timeline). The viral vector contained the neuron-specific human synapsin promoter (hSyn) for transfecting cortical neurons and carried along the humanized channelrhodopsin with the H134R mutation for optogenetic activation through cation conductance channelrhodopsin (hChR2(H134R)). To visualize the expression of the transgene in transfected cells the virus also contained the enhanced yellow fluorescent protein (eYFP).

After a surgical craniotomy over dorsomedial V1 performed under general anesthesia, we then proceeded to inject an average volume of 24 μl of the viral solution (VL: 25.5 μl, DP: 25 μl, AL: 22 μl, FL: 24.5 μl of AAV9-hSyn-ChR2-eYFP (UPenn Lot: CS0964 based on Addgene 26973P, titer: 1.03e13 GC/ml). The construct was loaded into a 10 μl Nanofil syringe (World Precision Instruments) to which a beveled 34 GA needle (World Precision Instruments) was assembled prior to injections into the cortex under microscopic control using a microinjection system (UMP3-1, SYS-Micro 4; World Precision Instruments). To increase optogenetic responses after one year, animal DP was reinjected at ∼ 4mm lateral and posterior of the first injection site using 28.5 μl of the same construct repackaged in serotype AAV5, which provided a higher titer (AAV5-hSyn-ChR2-eYFP, UPenn Lot: CS1078 based on Addgene 26973P, titer: 3.828e13 GC/ml). All injections were made in five different locations in a plus-shaped pattern within the chamber and separated by 2 mm horizontally and vertically (see example in **Supp. Fig. 1B** of monkey VL). On rare occasions the injection site was slightly modified to avoid the penetration of blood vessels. To improve viral expression across different cortical laminae, we injected the construct at three different depths starting with the deepest (approx. 1500µm) followed by a central (approx. 1000µm) and a superficial injection (approx 500µm) (see **Supp. Fig. 1C**). The intended injection depth was confirmed upon retraction by visualizing the exit of the injection needle from the cortex. In each depth, we injected a volume of approx. 1500 nl over the course of six minutes at a rate of 4 *nl* per second. To compensate for pressure induced by the injection needle, we allowed the tissue to settle for one to two minutes after having moved it to the different depths and also after each injection. Following the injection procedure, the craniotomy was covered with a custom-fit MRI compatible recording chamber (Ortiz-Rios et al., 2018).

#### Optical stimulation parameters for fMRI and electrophysiology

For the fMRI experiments, light (LED, Prizmatix UHP System) was delivered during 30s blocks followed by 30s blocks of no stimulation. For the ON block, light was either continuous or a train of square pulses for the duration of the block; pulse trains were tested at 5Hz, 10Hz, and 40Hz at 50% duty cycles. TTL signals were used to modulate the LED and the stimulation frequnecy parameters were used in all experiments. The power of the 451 nm LED was between 40-62 mW and 50 mW for the 626 nm LED.

For the electrophysiology experiments, the same LED systems were used for epidural stimulation in monkey VL. Light was delivered continuously or pulsed for the stimulation period (1000ms) with a randomised intertrial interval (ITI) (1000-2000ms) with no stimulation. The power of the 451 nm LED was between 52-56mW and that of the 628 nm LED was 50mW. In monkey FL, blue light (LuxX 473 nm diode laser, Omicron Lighthub-4) was delivered intracortically via an embedded optical fibre in the electrode array. For continuous stimulation conditions, stimulation duration was limited to 300 ms to avoid any potential damage. For pulsed stimulation, stimulation duration was 1000 ms with an ITI of 1000-2000 ms in all conditions. 473 nm laser power was between 37.5-55mW (mean = 48.8mW). Red light (594 nm DPSS laser, Omicron Lighthub-4) was only tested continuously with a stimulation period of 300 ms and power levels of ∼40mW.

#### MRI data acquisition

During the scanning period, animals remained calm in darkness during the opto-stimulation periods. Alternated runs with either red-light or movie watching were additionally performed. A vertical 4.7 Tesla magnet, running ParaVision 5.1 (Bruker, BioSpin GmbH, Ettlingen, Germany) and equipped with a 4-channel phase-array coil that covered the whole head (https://www.wkscientific.com), was used to acquire MR images. Imaging data consisted of two types of datasets: T1 for anatomical analyses and echo-planar imaging (EPI) for functional analyses (**Fig. 1**). We used the same parameters and hardware settings across all four subjects and experimental sessions.

For the first experiment in monkey VL.LV1 and one session in FL (FL.Ia1) we used a magnetization-prepared Rapid gradient echo (MP-RAGE) sequence to acquire anatomical (T1) images: FA (°) = 30; TE (ms) = 3.95; TR (ms) = 2000; TI (ms) = 750; Accel. Factor = 1; RIM-RO (px) = 176; RIM-PH (px) = 176; RIM-SL (px) = 72; RR-RO (mm) = 0.6; RR-PH (mm) = 0.6; RR-SL (mm) = 0.6; TA (h:m:s) = 0:14:48. For functional data on VL.LV1 and FL.Ia1 sessions we used a gradient-echo (GE) EPI sequence was used to acquire BOLD signal modulation for functional mapping with the following parameters: FA (°) = 65; TE (ms) = 21; TR (ms) = 1500; BW (Hz/Px) = 1704; ES (ms) = 58; Accel. Factor = 2; RR-RO (mm) = 1.2; RR-PH (mm) = 1.2; RR-SL (mm) = 1.2; RIM-RO (px) = 88; RIM-PH (px) = 88; RIM-SL (px) = 31; Nacq = 200; TA (h:m:s) = 0:5:00.

For all subsequent sessions we increased the resolution to obtain more detailed maps of the local activation region. For all monkeys and sessions (VL.MQ1, VL.M31, DP.NJ1, DP.Qq1, AL.SD1, AL.SI1, FL.Uv1, FL.Uy1) we used the following anatomical and functional parameters: MP-RAGE sequence for anatomical images: FA (°) = 30; TE (ms) = 5.42; TR (ms) = 2000; TI (ms) = 750; Accel. Factor = 1; RIM-RO (px) = 70; RIM-PH (px) = 82; RIM-SL (px) = 26; RR-RO (mm) = 0.27; RR-PH (mm) = 0.25; RR-SL (mm) = 1.4; TA (h:m:s) = 0:20:48. For functional data we used a gradient-echo (GE) EPI sequence with the following parameters: FA (°) = 65; TE (ms) = 21; TR (ms) = 1700; BW (Hz/Px) = 3409; ES (ms) = 58; Accel. Factor = 2.3; RR-RO (mm) = 0.79; RR-PH (mm) = 0.79; RR-SL (mm) = 1.4; RIM-RO (px) = 70; RIM-PH (px) = 70; RIM-SL (px) = 20; Nacq = 176; TA (h:m:s) = 0:5:00.

#### fMRI data analyses

All pre-processing analyses were conducted using open software provided in the packages of AFNI (Cox, 1996), SUMA (Saad, Reynolds, Argall, Japee, & Cox, 2004). For anatomical scans, the D99 atlas (Reveley et al., 2017) was used to re-create an in-session anatomical surrogate brain of each monkey. White matter and grey matter were obtained from the in-session atlas ROIs parcellation of the warped D99 template. The resulting masks were then used to render a white, pial, and flattened surface using Freesurfer (Dale, Fischl, & Sereno, 1999). For displaying the ROIs into the surface, we first created a color mappable niml file of the atlas volume using the AFNI program (3dVol2Surf). In SUMA, we then display the contours of each ROI on the surface along with the activation maps.

Pre-processing of fMRI time series followed standard preprocessing steps which included: slice-timing correction, motion correction, spatial-smoothing (3dmerge, 1.5 mm fwhm), and mean scaling of the time series. 3dDeconvolve was used for linear least-squares detrending to remove nonspecific variations (i.e., scanner drift) and regression that included the overall white matter resulting in a residual time-series which were then used for coherence analyses.

Coherence analyses were then performed on the residual time series by measuring the ratio between the amplitude at the fundamental frequency to the signal variance, ranging between 0 and 1 (Brewer, Press, Logothetis, & Wandell, 2002). The measure of coherence is

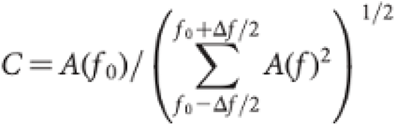

where is the stimulus frequency, the amplitude of the signal at that frequency, the amplitude of the harmonic term at the voxel temporal frequency, and the bandwidth of frequencies in cycles/scan around the fundamental frequency. For all opto stimulation, it corresponds to one cycle (1/60 sec = 0.016 Hz) and corresponds to the frequencies around the fundamental (see **Fig. 1D**). The threshold was chosen at a coherence level > 0.35.

#### Electrophysiological recordings

In both monkeys, the activity of all the cortical layers was recorded with laminar probes. In monkey VL, recordings were performed using dual-shaft laminar Atlas probes (Atlas Neuro, Leuven, Belgium) with 16 electrodes in each shaft (150µm electrode spacing, 50µm electrode diameter). An optic fibre (1.5 mm diameter, 0.5 N.A.) was placed outside the cortex to deliver light (451 nm; blue light for stimulation, and 626 nm; red light for control) from an external LED system (Prizmatix UHP System).

In monkey FL, neurophysiological data was acquired using a laminar Plexon S-probe (Plexon Inc., Texas, USA) with 24 contacts (100µm electrode spacing, 15µm electrode diameter), a total length of 100mm and a probe diameter of 300 µm. The S probe also had a fiber optic channel installed between electrodes 8 and 9 allowing for intracortical optogenetic stimulation. We used an Omicron Laser System (Omicron Lighthub-4) both for stimulation (LuxX 473 nm diode laser, nominal aperture, ∼50mW) and control (594 nm DPSS laser, nominal aperture, ∼40mW).

Mock laser and LED stimulation were used to assess for neuronal responses unrelated to optogenetic activation but to artifacts induced by temperature or non-wavelength specific light modulations. Triggers to the Laser and LED systems were sent and recorded using a behavioural experiment software (MWorks) via an interface card (National Instruments).

#### Analysis of electrophysiological signals

Raw electrophysiological signals were recorded using a Blackock recording system (BlackRock Microsystems, Inc.) and sampled at 30kS/s. Data analysis was done in Matlab using the Fieldtrip toolbox (Oostenveld et al., 2011) and custom scripts. The CSDplotter toolbox (Pettersen et al., 2006) was used to calculate the CSD profiles of the visual responses using the standard CSD method. LFP signals were obtained by downsampling the raw signal to 500Hz and then applying a notch filter at 50Hz to remove any line noise. Multiunit activity signals were obtained by applying a high pass filter at 250 Hz to the raw signal and then units were extracted using a threshold that is 2.5-3.5 std of the channel average. MUAe (multiunit activity envelope) signals were obtained by applying a high pass filter at 300 Hz to the raw signal, rectifying and then downsampling the signal to 500Hz (Supèr and Roelfsema, 2004). Optogenetic stimulation trials were extracted based on the onset of the TTL signal sent to the laser by the NI card. In case of a pulse train, the first pulse onset is used as a marker. The first 100ms after stimulation onset were excluded from the laminar activity pattern calculation to avoid the transient response and any artifacts caused by the onset of the light.

For epidural stimulation, light onset artifacts were present. They could be removed manually during multiunit activity extraction, however it was not possible in MUAe. For intracortical stimulation, however, no light artifacts were observed in the MUA/MUAe signals. Response onset for visual and optogenetically modulated responses were calculated by detecting when the signal crosses 4*std of the baseline.

The laminar activation pattern was calculated using the sustained (100-300ms post stimulation onset) multiunit activity. For each channel in a session, the trials were averaged and the mean the sustained period was calculated. The laminar activation pattern was calculated for each session and then normalised by the maximum firing rate for that session.

#### Two-alternative-choice (2-AC) behavioral task

Animals were positioned at a distance of 84cm in front of a ViewPIXX computer screen (RR: 120Hz, VPIXX technologies), allowing for the presentation of stimuli on the screen at +/- 16° horizontally and +/- 9.5° vertically. Behavioural tasks were programmed and executed using MWorks (MWorks) with custom scripts that used the MWorks Experiment Language (MWEL).

The animals’ eye movements were calibrated to the eye tracking system such that we were able to infer their fixation location from a stimulus position on the screen and vice versa. During visual calibration animals were performing saccades to visual stimuli that appeared at different locations (12 points) on the screen. The animals were trained to fixate on these stimuli for 500 ms with adequate precision (<2° visual angle). Upon successful fixation the animals received a reward in the form of fruit juice or water. Once calibrated animals were performing receptive field (RF) mapping and visual detection tasks.

Automatic RF mapping was performed in order to identify the spatial location of the minimum RF of underlying neuronal populations. To do so, animals were required to maintain precise central fixation (1°, 1000 ms). During central fixation, black squares (diameter: 0.5/1°) were presented on a 5×5 grid in the lower left quadrant of the visual field (duration: 100/grid location), where the RF was located, while electrophysiological activity was recorded. The RF was estimated based on MUAe at each grid location.

Subsequently, animals were required to perform a two alternative forced choice task (2AFC) with three equally probable conditions. During all tasks, the animals needed to maintain precise fixation on a 0.2° central fixation point (within a window <1°, for 500 - 1250 ms). During central fixation, two targets (0.3° red discs) appeared in the upper left and upper right part of the screen. Fixation time was followed by (1) the presentation of a 1° white disc with varying contrasts on the left side of the central fixation point inside the RF that was previously determined, (2) no visual or optical stimuli (control condition) or (3) continuous optogenetic stimulation. The animal was initially trained a version of the task that does not have optogenetic stimulation; upon detection of the disc on the left of the central fixation point, the animals were trained to perform saccades to the left target indicating a correct detection trial and to the right target, if otherwise, indicating a correct catch trial. Later on, in addition to visual and catch trials, neurons were stimulated optogenetically, thus potentially inducing a phosphene in the RF. The animals received a reward if they performed a saccade to the left in the opto trials.

We used d’ as a measure of sensitivity. d’ is the difference between the z-transforms of hits and false alarms. A value of d’ was calculated for the optogenetic condition and every visual condition. Catch trials were used to estimate false alarms and correct rejections.

#### Eye movement analyses

Eye position was recorded with Eyelink 1000 plus with a sampling rate of 1000 Hz; the raw signal was calibrated and converted to visual degrees by Mworks. We detected eye-movements automatically with a usually employed velocity based algorithm (see (Engbert and Kliegl, 2003), for details). Velocity was calculated using a 5ms moving window. An eye movement was classified as a saccade if velocity increased beyond 6 std. Reaction times were calculated as the time interval between the onet of the stimulus and the onset of the saccade. No latencies were below 100 ms.

#### Histological analysis of brain slices

Once sufficient experimental data was gathered, animals were anaesthetised with an overdose of anesthetic and then perfused through the heart with phosphate-buffered saline (37°, pH 7.4), followed by paraformaldehyde (40%, pH 7.4). The brains were then carefully removed and placed in Paraformaldehyde (40%, pH 7.4) over night. Subsequently, the brain was placed in increasing sucrose concentrations (10%, 20%, 30%) for cryoprotection.

Tissue was then cut on a freezing microtome (50 µm sections) and then stained using Cresyl violet before being mounted on microscopy slides. A subset of sections was also stained using a standard immunohistological protocol. Sections were incubated in a trisodium citrate buffer (pH 6, 0.1M, 1% horse serum, 1% goat serum, 0.3% TritonX) for 60 min. Sections were then incubated with a primary antibody for the vesicular glutamate transporter (vGlut2, MAB5504, Merckmillipore) used to label cells in the cortical layer V1 4C beta (Brodmann’s nomenclature), as has been shown previously (Balaram et al. 2014), diluted in PBS for 24 hours, before being treated with AlexaFluor conjugated secondary antibodies for two hours. Sections were then mounted on frosted microscope slides, covered with mounting medium conjugated with a nuclear cell body marker (Fluoroshield TM with DAPI, Merck) and protected using standard coverglass. Following treatment, representative sections were imaged using a fluorescence microscope (DM6B, Leica Navigator).

## Figure Legends

**Supplementary Figure 1.**
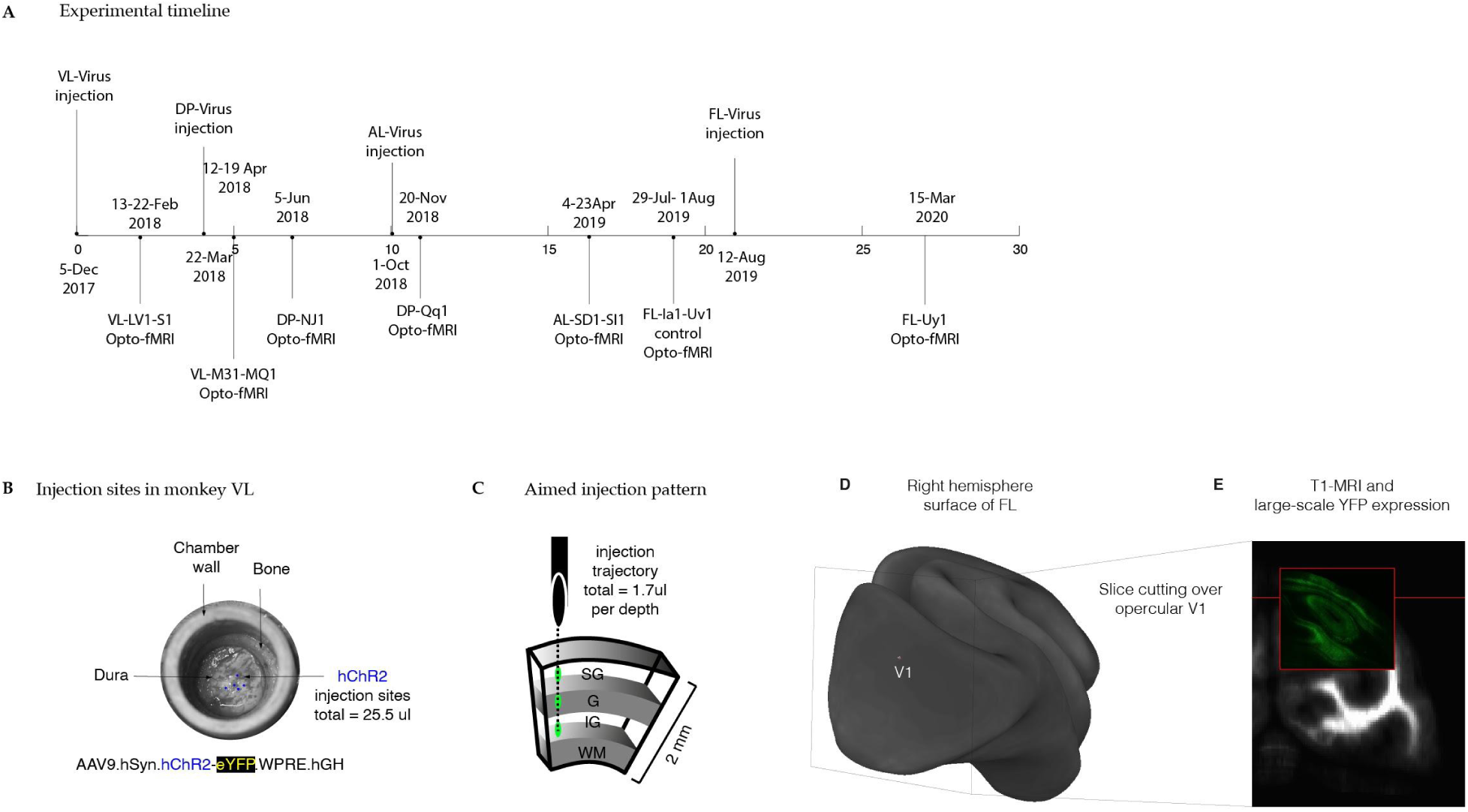
Experimental timeline and injection approach. **A**. Experimental timeline showing events for injections and opto-fMRI experimental sessions lasting 33 months. **B**. Photograph of chamber region with injection sites in monkey VL showing the cross-shaped injection pattern into cortical tissue. **C**. Illustration of the injection approach for each site with virus delivery at three different cortical depths with a total of 25.5*μl* of viral solution. **D**. Illustration of the V1 position of the slice showing eYFP expression in Figure 2B.

**Supplementary Figure 2.**
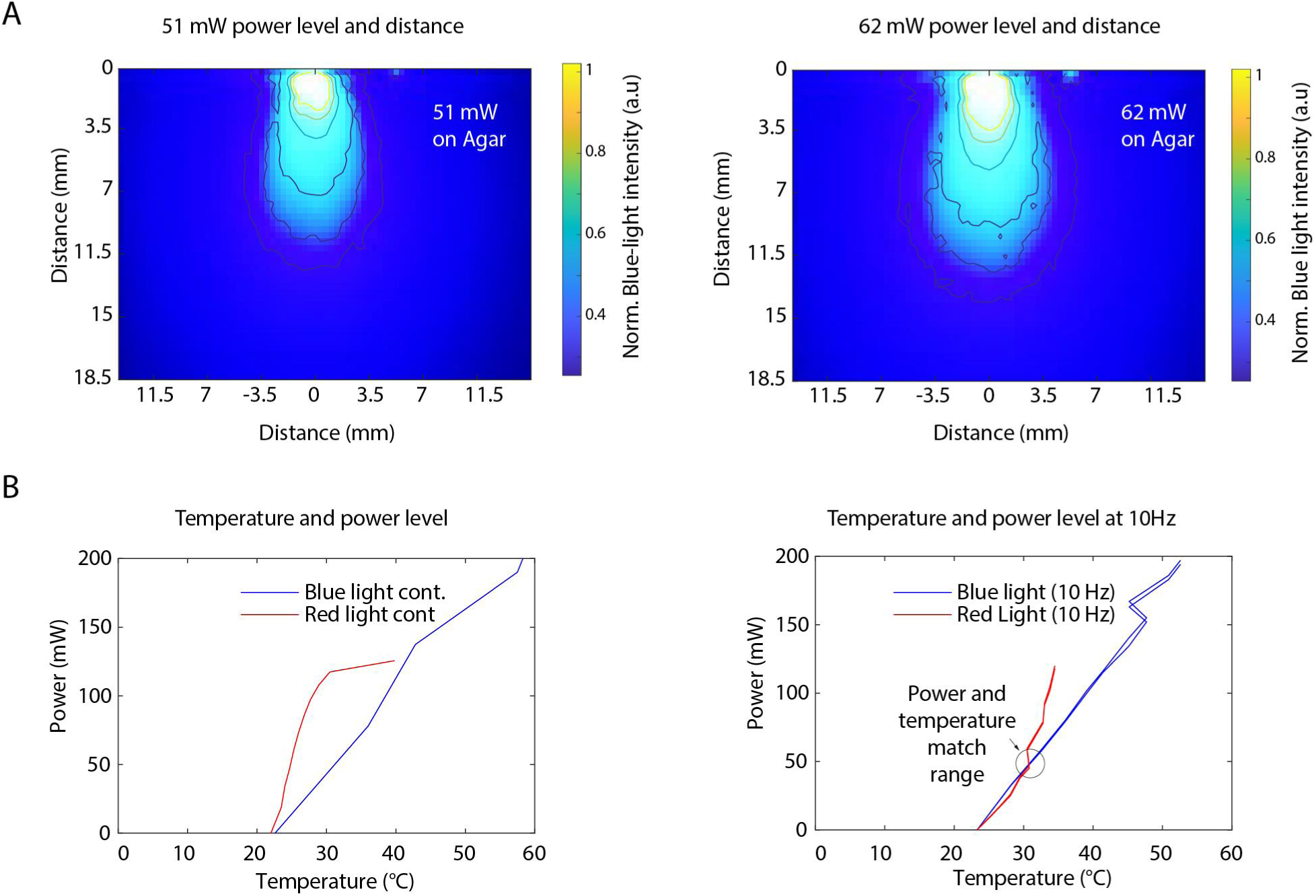
Power level, distance and temperature control measures in agar. **A**. Shows blue-light light emission within agar solution to mimic spread in the brain The light was delivered at two power levels 51 mW and 62 mW. The normalized light intensity shows the increase spread at higher powers with most light intensity more effective near the tip of the LED fiber optic and with a decay as a function of distance. **B**. Increase in temperature from 30s long blue-light vs red-light stimulation during continuous (left plot) vs 10 Hz modulation (right plot). Red-light reaches a temperature plateau before blue-light during continuous stimulation.

**Supplementary Figure 3.**
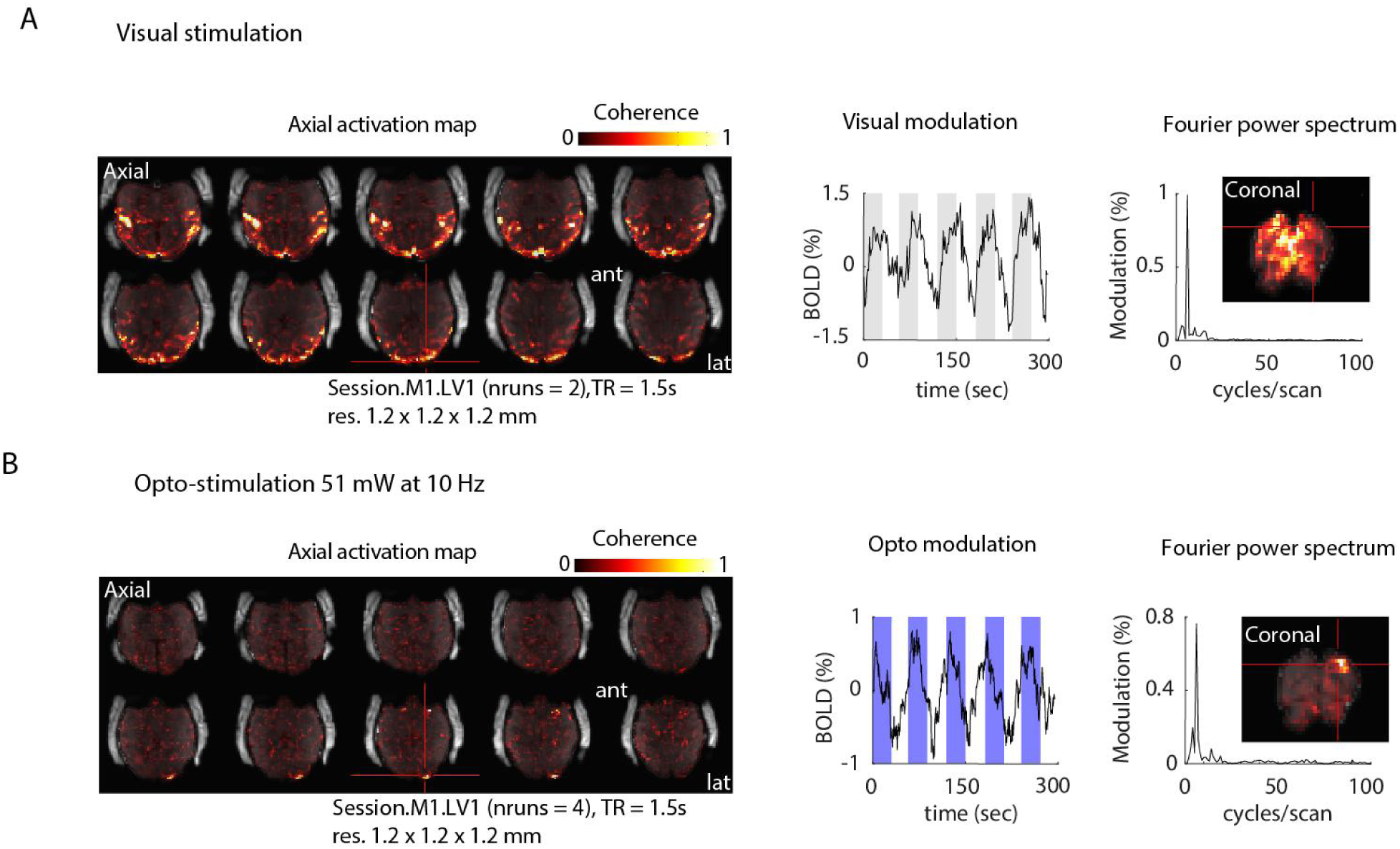
Comparison of V1 BOLD fMRI activation patterns elicited by optogenetic stimulation versus free-viewing of movie sequences. **A**. Overall coherence map without threshold shows regions with significant BOLD activation in V1 and extrastriate cortex elicited by the free-viewing of natural scene movies. The middle panel shows the time course of BOLD signal modulation from an exemplary voxel in the recording chamber region. The right side panel shows the associated power spectrum with a peak at the stimulation frequency (0.016 Hz, 1/60 sec). The inset shows the coronal slice of the activation in the visual cortex. **B**. A similar plot as in A for the Opto-fMRI stimulation shows BOLD activation in response to optogenetic stimulation at 10Hz. Note that elicited activation is restricted to V1. Middle and right panels show BOLD signal modulation and power spectrum of the same V1 voxel with a modulation rate peak of 0.016 Hz or 1 cycle/60 sec.

**Supplementary Figure 4.**
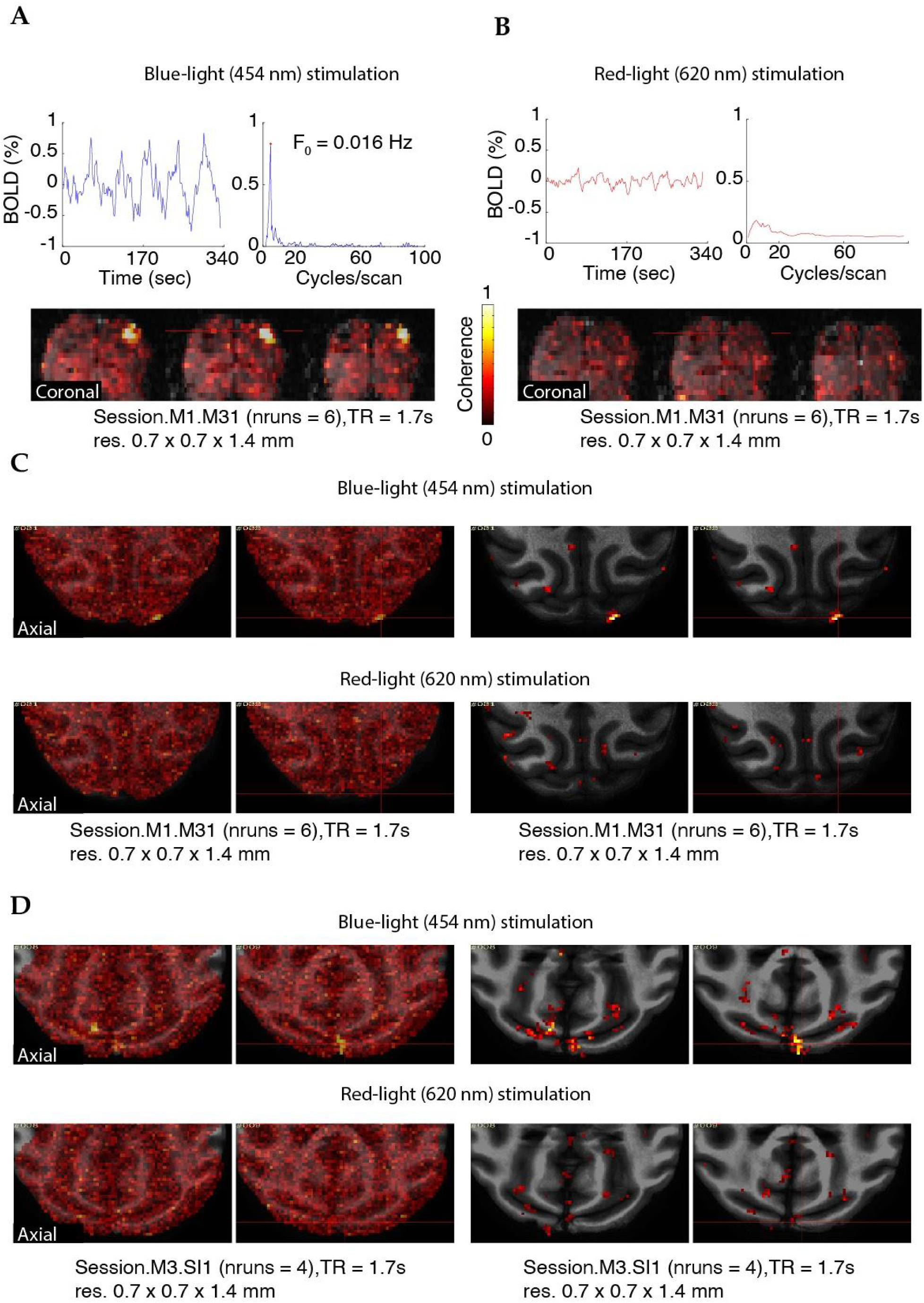
Wavelength specific stimulation drives the local positive BOLD response only after blue but not red-light stimulation. **A**. Example voxel time course and power spectrum of BOLD signal modulation from blue-light (451 nm, 50 mW at 10Hz). Bottom panel shows the coherence map without a threshold of three coronal slices (n runs = 4) in monkey VL. **B**. Same voxel time course and power spectrum of BOLD signal modulation after red-light stimulation (626 nm, 50 mW at 10Hz). Note the absence of BOLD modulation under this stimulation condition. **C**. Shows the same coherence map as in B for VL in the axial plane for both blue and red-light conditions. see coherence map without threshold. **D**. The coherence map without threshold for monkey AL in the axial plane for both blue and red-light conditions.

**Supplementary Figure 5.**
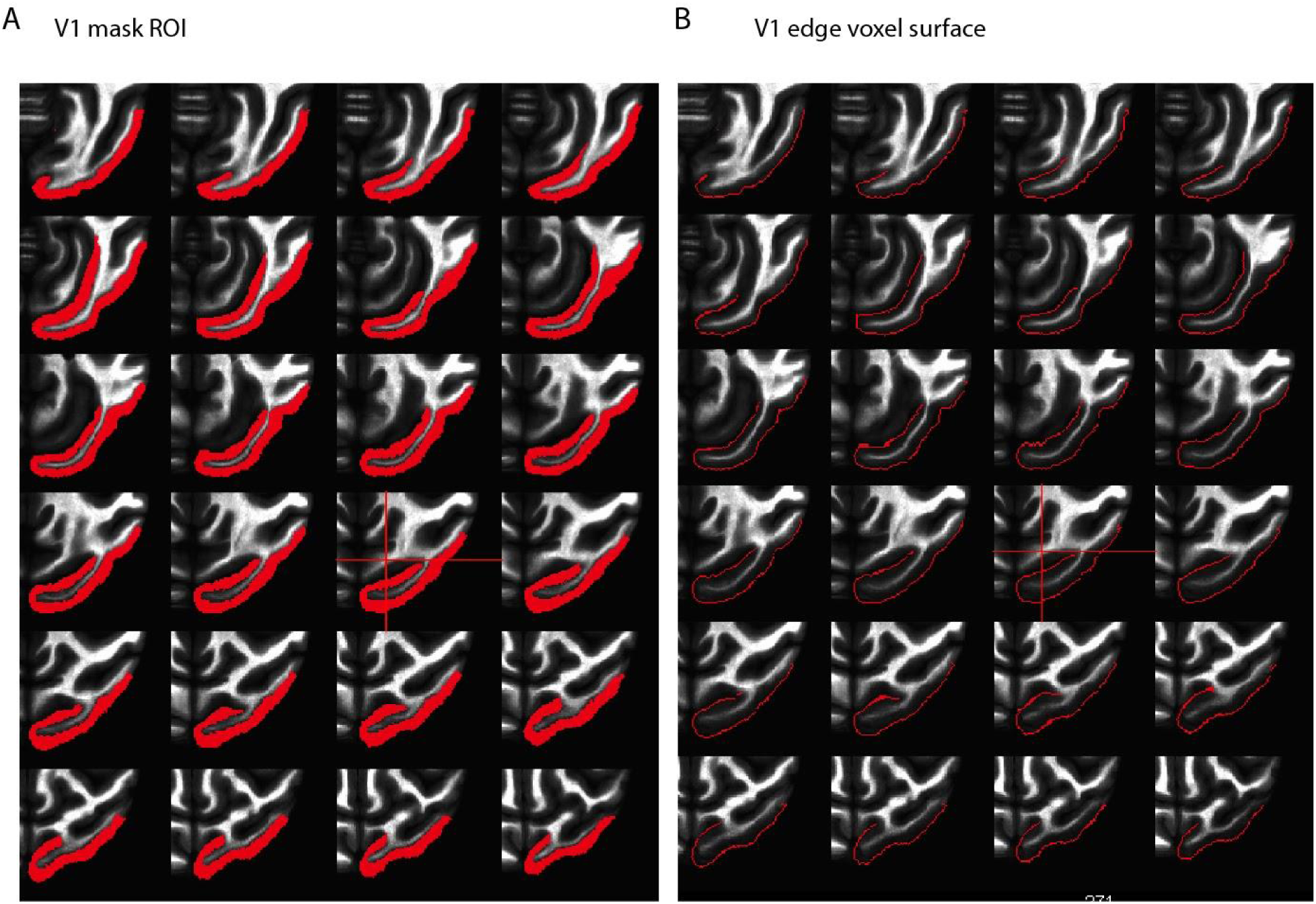
Region of interest ROI overlaid on the anatomy of V1. **A.** V1 cortex of monkey DP on an axial slice. **B**. V1 edge voxels of monkey DP on an axial slice. Both samples were used to quantify the extent of local BOLD response.

**Supplementary Figure 6.**
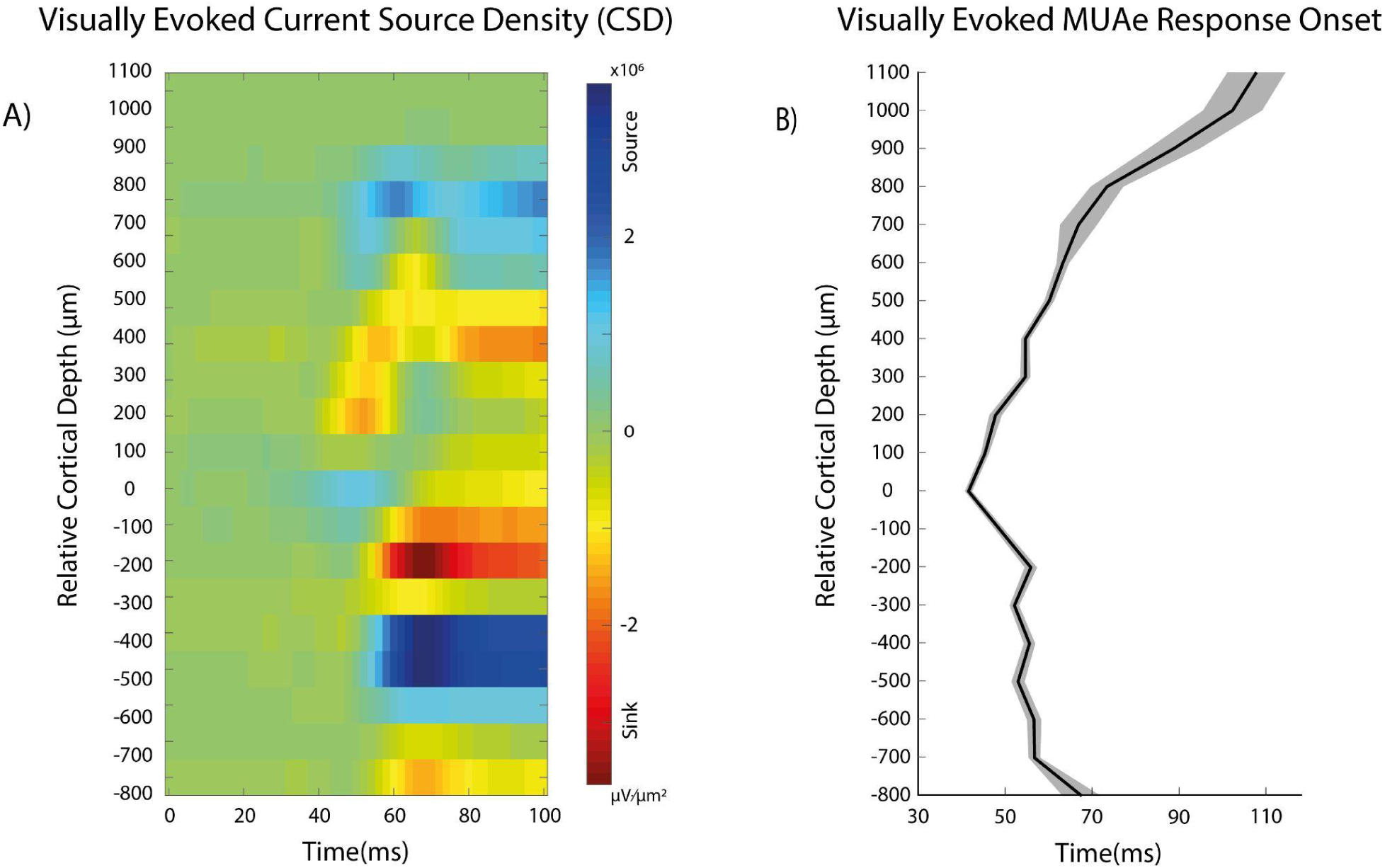
Current Source Density (CSD) and latency of visually evoked MUAe responses in V1 as a function of relative cortical depth. **A.** Example CSD response profile to a 5° patch of drifting gratings (spatial frequency 2 cycles/°, speed: 2-4 cycles/s) from one session. The lower border of the first sink (in red, thalamic input to L4C) is used as a reference to align laminar data across sessions. **B.** Response onset latency for the visually evoked MUAe across cortical depth across sessions. Latency onset was extracted when the signal crosses 4*SD of the baseline.

**Supplementary Figure 7.**
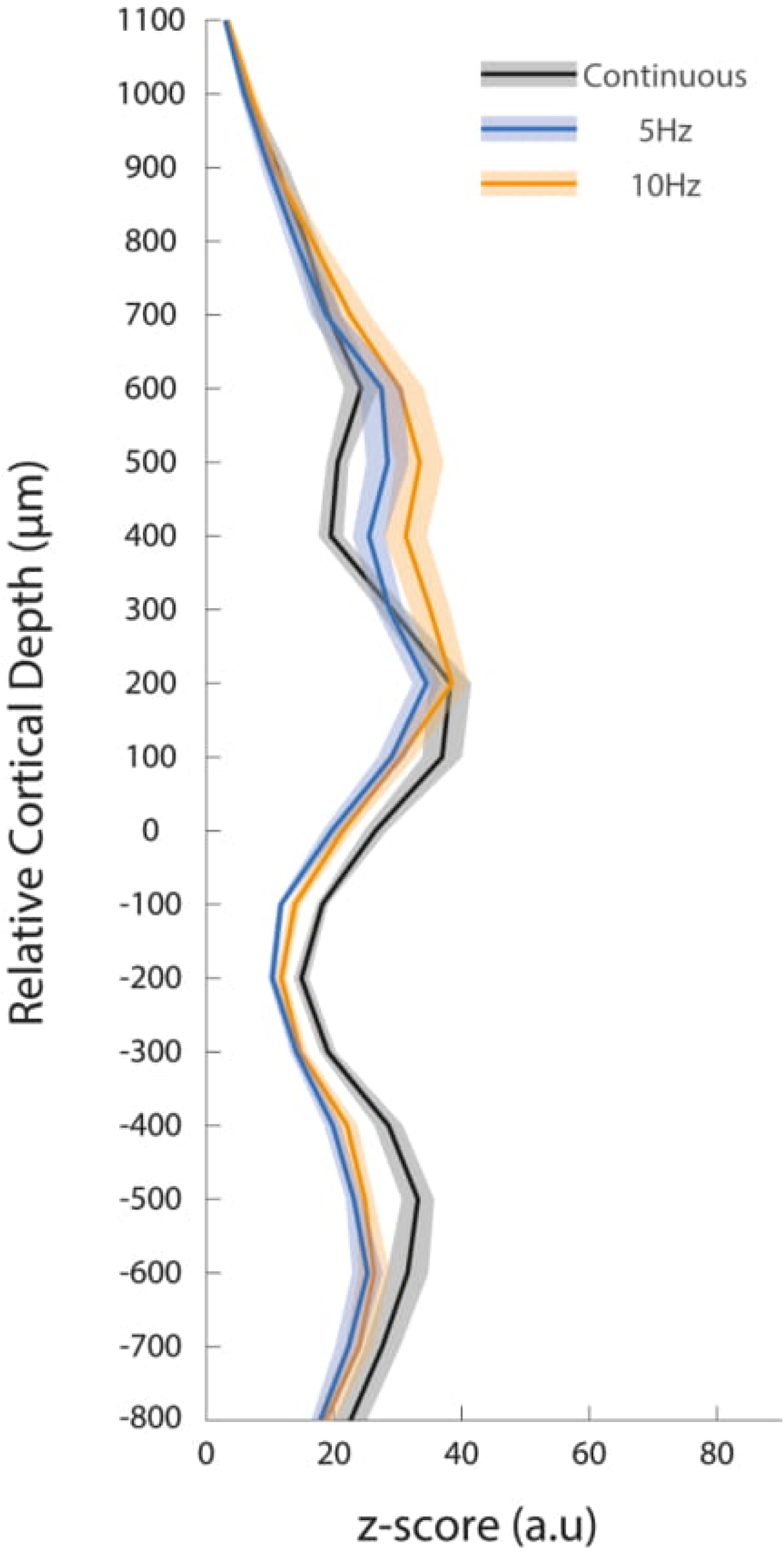
Laminar activation pattern for optogenetic stimulation frequencies 5Hz and 10Hz. Similar to Fig. 3E, the laminar activation pattern is calculated for continuous (black) stimulation as well as 5Hz (blue) and 10Hz (orange) stimulation frequencies. The average MUEe firing rates are calculated for 100-300ms post stimulation onset for the continuous stimulation and 100-900ms post the first stimulation pulse onset for the pulsed stimulation.

**Supplementary Figure 8.**
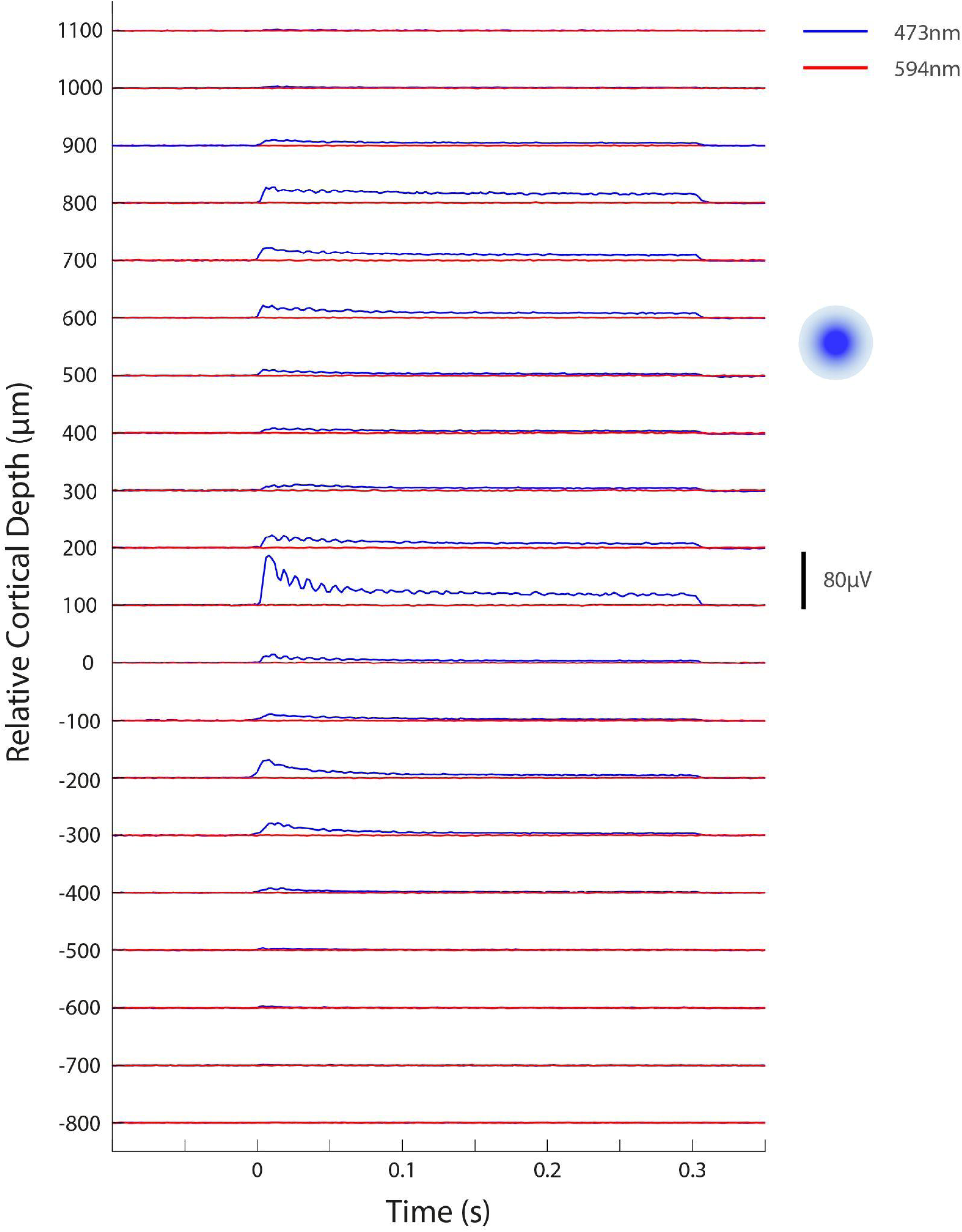
Example MUAe responses to both continuous blue (473nm) and continuous red (594nm) laser stimulation. Red light stimulation (40mW) did not have an effect on neural activity (red) when compared to blue light stimulation with a similar power (37.8mW). The location of the embedded fibre is indicated by the blue circle on the right.

**Supplementary Figure 9.**
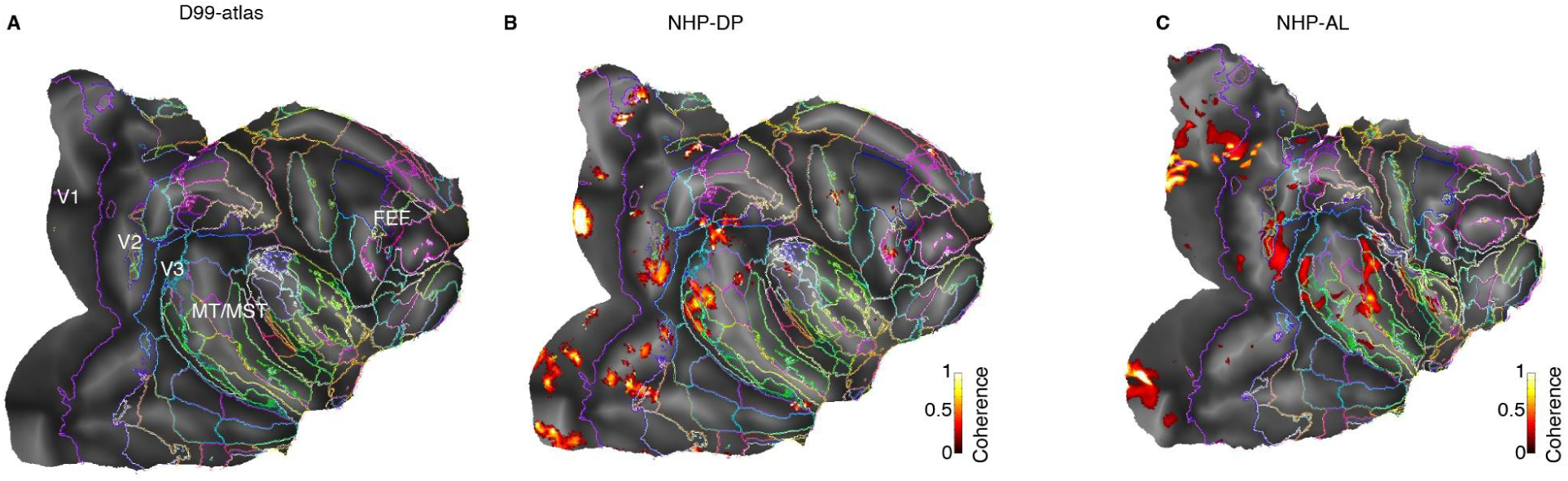
V1 optogenetic stimulation drives BOLD activation in higher cortical areas. **A**. Flat map of D99-atlas showing the overall cortical parcellation for the average brian map. Activation maps showing extrastriate BOLD activity in V1, V2, V3 and motion complex regions MT/MST in NHPs DP (**B**) and AL (**C**).

**Supplementary Figure 10.**
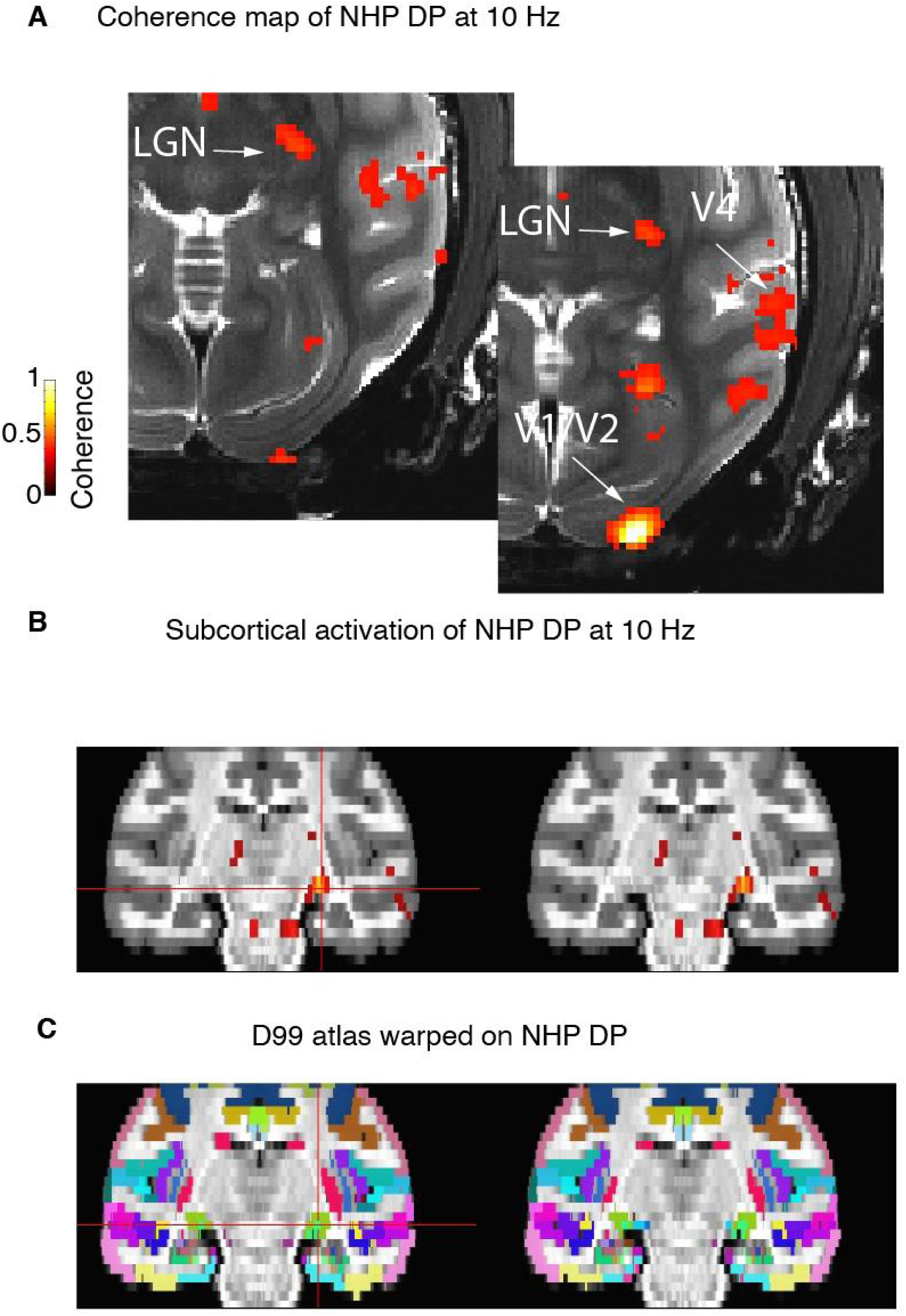
Extrastriate and subcortical activation maps in NHP DP. Similar to Fig. 3E, **F**. Two axial slices of monkey DP showing significantly active regions V1, V2 V4 and LGN. **B**. Two coronal slices outline the lateral geniculate nucleus (LGN) which show significant activation in the stimulated hemisphere. **C**. Cortical regions form the D99 atlas mapped onto the warped template of monkey DP.

## Acknowledgments

We wish to thank Ian Milne and Joe Wardle for their help in developing hardware and equipment for MRI-compatible imaging. We would also like to thank the CBC staff for their constant effort with animal training and handling during experiments. We received advice from Carsten Klein, Eric Schmiedlin and Henri Evrard on optogenetic procedures and histology for which we are very grateful. We would also like to thank Midea Ortiz-Rios, Christine Roulin and Veronique Moret for their help in preparing the brain tissue. We are grateful to the constant support, discussion and encouragement by Alex Thiele and Chris Petkov and to numerous fun joint lab meetings We thank Alex Thiele, Chris Petkov and Pavel Kusmierek for comments on this manuscript. This work was supported by ERC OptoVision 637638 granted to Michael C. Schmid.

